# The EBNA2-EBF1 complex promotes oncogenic MYC expression levels and metabolic processes required for cell cycle progression of Epstein-Barr virus-infected B cells

**DOI:** 10.1101/2021.12.29.474426

**Authors:** Sophie Beer, Lucas E. Wange, Xiang Zhang, Cornelia Kuklik-Roos, Wolfgang Enard, Wolfgang Hammerschmidt, Antonio Scialdone, Bettina Kempkes

## Abstract

Epstein-Barr virus (EBV) is a human tumor virus, which preferentially infects resting human B cells. Upon infection *in vitro*, EBV activates and immortalizes these cells. The viral latent protein EBV nuclear antigen (EBNA) 2 is essential for B cell activation and immortalization; it targets and binds the cellular and ubiquitously expressed DNA binding protein CBF1, thereby transactivating a plethora of viral and cellular genes. In addition, EBNA2 uses its N-terminal dimerization (END) domain to bind early B cell factor (EBF) 1, a pioneer transcription factor specifying the B cell lineage. We found that EBNA2 exploits EBF1 to support key metabolic processes and to foster cell cycle progression of infected B cells in their first cell cycles upon activation. An α1-helix within the END domain was found to promote EBF1 binding. EBV mutants lacking the α1-helix in EBNA2 can infect and activate B cells efficiently, but the activated cells fail to complete the early S phase of their initial cell cycle. Expression of *MYC*, target genes of MYC and E2F as well as multiple metabolic processes linked to cell cycle progression are impaired in EBVΔα1 infected B cells. Our findings indicate that EBF1 controls B cell activation via EBNA2 and, thus, has a critical role in regulating the cell cycle of EBV infected B cells. This is a function of EBF1 going beyond its well-known contribution to B cell lineage specification.

**Significance statement:** Epstein-Barr virus (EBV) infects primary B cells and establishes life-long latent infection in these cells. EBV nuclear antigen (EBNA) 2 drives early processes of B cell activation and cell cycle entry. The surface of the N-terminal dimerization domain of EBNA2 exposes a five amino acid α-helix (α1) that recruits EBF1 to activate *MYC* and downstream targets of both MYC and E2F to support critical metabolic processes in infected B cells and to drive them through S phase in the first cell cycle post-infection. Our study demonstrates how EBNA2 exploits EBF1, a key factor of B cell lineage specification to initiate proliferation and high-lights the α1-helix as a potential Achilles heel of the virus at the stage when latent infection is established.

## INTRODUCTION

Epstein Barr virus (EBV) is a γ-herpesvirus, which is associated with diverse malignancies such as nasopharyngeal- and gastric carcinoma, post-transplant lymphoproliferative disease, Burkitt’s lymphoma, diffuse large B cell, Hodgkin’s and NK/T cell lymphoma (Shannon-Lowe, Rickinson and Bell, 2017). Like other herpesviruses, EBV switches between lytic infection to produce infectious virus and latent infection to maintain a lifelong persistent reservoir in the infected host. EBV can infect epithelial and lymphoid cells but the viral reservoir of EBV *in vivo* are non-proliferating memory B cells. To reach this long-lived B cell compartment, the virus initially infects resting human B cells, activates these and drives their proliferation. This process is considered to be the essential step to initiate persistent infection *in vivo*, can be studied in short-term and long-term cell culture infection models and is referred to as B cell immortalization (Münz, 2019). It requires the timely expression and collaborative action of a group of nuclear viral proteins termed EBNA1, 2, 3A, 3B, 3C and EBNA-LP as well as a group of latent membrane proteins termed LMP1 and LMP2.

EBNA2 initiates the immortalization process by directly and immediately activating cellular target genes including the cellular proto-oncogene *MYC* (Kaiser *et al*., 1999) as well as by activating certain viral promoters. Several studies have demonstrated that mutant EBVs devoid of EBNA2 are incapable to immortalize primary human B cells (Cohen *et al*., 1989; Hammerschmidt and Sugden, 1989; Cohen, Wang and Kieff, 1991; Kempkes, Pich, *et al*., 1995; Kempkes, Spitkovsky, *et al*., 1995). Most recently, a panel of viral mutants deficient for the expression of single latent genes has been studied during the early phase of the immortalization process. Apart from EBV devoid of EBNA2, all EBV mutants efficiently supported cell cycle activation. This finding highlights the essential role of EBNA2 to initiate the complex process of B cell immortalization (Pich *et al*., 2019).

EBNA2 is a transcription factor that uses cellular sequence-specific DNA binding proteins as DNA anchors since it does not bind DNA directly. EBNA2’s long-known anchor protein is CBF1, a sequence specific DNA binding factor, which is highly conserved and ubiquitously expressed. EBNA2 mutants incapable of CBF1 binding fail to immortalize B cells *in vitro* (Yalamanchili *et al*., 1994). Chromatin immunoprecipitations (ChIP) studies map EBNA2 binding to B cell specific super enhancers, which are present in primary resting B cells prior to infection (Zhao *et al*., 2011; Jiang *et al*., 2017). Due to the ubiquitous expression of CBF1 in all human cells, it does not confer B cell specificity to EBNA2. Early B cell factor (EBF) 1, which is expressed exclusively in B cells, is another DNA binding protein associated with EBNA2 (Glaser *et al*., 2017). Interestingly, EBF1 and CBF1 frequently co-occupy high affinity EBNA2 binding sites and EBNA2 can enhance EBF1 and CBF1 signal intensities demonstrated by ChIP sequencing experiments. This indicates that EBNA2 not only uses transcription factors as DNA anchors but also influences transcription factor densities at enhancer and super-enhancer sites in B cells (Zhou *et al*., 2015; Lu *et al*., 2016). The EBNA2 N-terminal dimerization (END) domain forms a dimeric globular structure (Friberg *et al*., 2015) and is sufficient to mediate the interaction between EBNA2 and EBF1. Furthermore, we found that EBF1 promotes the assembly of EBNA2 chromatin complexes in B cells (Glaser *et al*., 2017).

The pioneer factor EBF1 can open chromatin and control B cell specific gene expression and triggers *MYC* expression and proliferation during mouse early B cell development (Boller, Li and Grosschedl, 2018; Sigvardsson, 2018). In addition, EBF1 is required for the generation and survival of distinct mature mouse B cell populations and their proliferation in response to activating signals (Györy *et al*., 2012; Vilagos *et al*., 2012). To date, the function of EBF1 in primary human B cells has not been studied. In human cell lines, EBF1 expression strongly correlates with transcription of hallmark B cell genes confirming that EBF1 defines B cell identity in the human system as well. Whether EBF1 controls B cell proliferation could not be studied in the respective cell culture model systems (Bohle *et al*., 2013; Bullerwell *et al*., 2021).

Previously, we have presented the three-dimensional structure of the END domain. The 58 amino acid residue END domain forms a compact homodimer, which is stabilized by a hydrophobic interface between the two monomers. While interface mutants impaired dimerization, mutagenesis of selected amino acid residues exposed on the hydrophilic surface, H15 and the α1-helix, impaired EBNA2 transactivation but did not affect dimerization (Friberg *et al*., 2015). The goal of this study was to characterize the specific functional impact of EBF1 on the activity of EBNA2 by reverse genetics. Here, we show that an EBNA2 α1-helix deletion mutant (EBNA2Δα1) does not form complexes with EBF1 but bound perfectly well to chromatin regions known to tether EBNA2 via CBF1. EBV mutants carrying the α1-helix deletion in EBNA2 (EBVΔα1) infected and activated primary B cells, but cell cycle progression was severely compromised, and most cells arrested in the early S phase. Long-term cultures of EBVΔα1 infected B cells could only be expanded by continuous CD40 stimulation indicating that EBF1 is absolutely required to maintain B cells in their EBV-driven immortalized state. RNA-sequencing experiments revealed that EBVΔα1 poorly activated *MYC* and gene sets known to be targets of MYC and E2F and failed to promote metabolic processes linked to cell cycle progression and, thus, explained the failure of EBVΔα1 to immortalize primary B cells.

## RESULTS

### EBNA2-EBF1 complex formation requires the END domain with its α1-helix

To identify critical regions within the EBNA2 N-terminal dimerization (END) domain, we investigated mutants of the END domain in GST pulldown assays with EBF1 containing whole protein lysates. The single amino acid mutant H15A and a deletion of the α1-helix (E2Δα1) were analyzed (Figure 1B). While the H15A missense mutant retained most of its EBF1 binding, the END domain mutant lacking the α1-helix (E2Δα1) failed to bind EBF1 (Figure 1C). Next, EBNA2 mutants lacking the entire END domain (E2ΔEND), E2-H15A or E2Δα1 were co-expressed in a B cell line (DG75) together with EBF1 and tested for co-immunoprecipitation (Co-IP) using full-length EBNA2 as control. Data shown in panel D of figure 1 confirmed the results obtained by GST pulldown.

**Figure 1.**
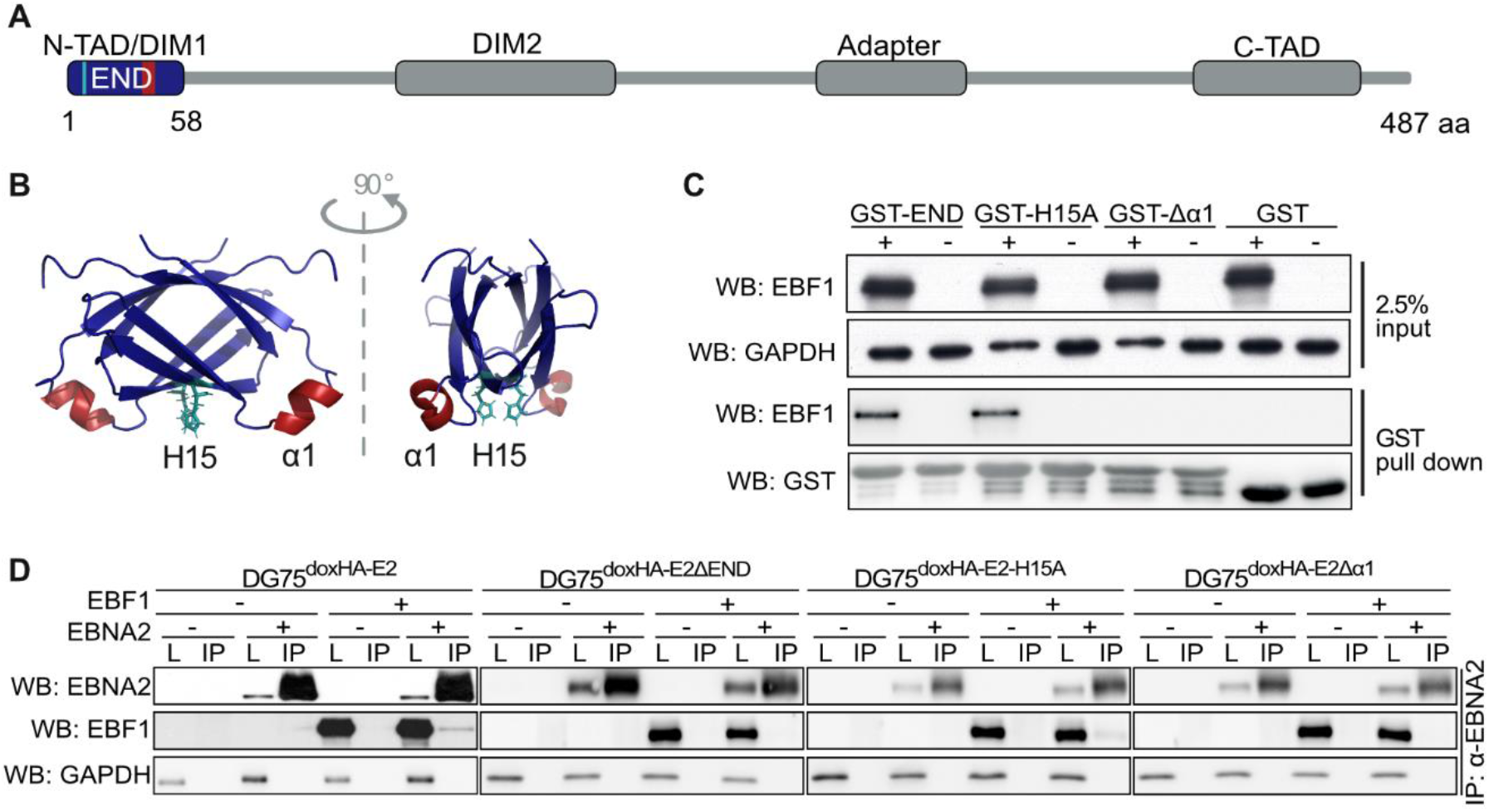
The α1-helix on the surface of the EBNA2 N-terminal dimerization domain (END) binds EBF1. (A) Schematic representation of functional EBNA2 modules: The END domain (blue) with positions of the α1-helix (aa 35-39; red) and histidine (H) 15 (turquoise), N- and C-terminal transactivation domains (N- /C-TAD), two dimerization domains (DIM1/2) and an adaptor region that confers DNA binding via the CBF1 protein. (B) NMR structure of the END domain homodimer, H15 (turquoise), α1-helix (red). (C) GST-pulldown experiments with the END domain and EBF1. Whole cell lysates of DG75 cells transfected with an EBF1 expression plasmid (+) or an empty vector control (-) were incubated with GST-END fusion proteins purified from E.coli: wild type END (GST-END), END-H15A missense mutant (GST-H15A), END α1-helix deletion mutant proteins (GST-Δα1) or GST. GAPDH served as a loading control for input lysates. (D) Co-IPs of HA-tagged EBNA2 (HA-E2) and EBNA2 mutants (HA-E2ΔEND, HA-E2-H15A or HA-E2Δα1) conditionally expressed in DG75 cells which were transiently transfected to express EBF1. Transfected EBF1 (+) or empty vector control (-), whole cell lysates (L), immunoprecipitation (IP). Immunoprecipitations were performed with EBNA2 specific antibodies. GAPDH served as loading control for total cell lysates.

### EBVΔα1 activates primary B cells but does not cause long-term proliferation of infected cells

The immortalization of B cells is a complex multi-step process. The key function of EBNA2 in the infected B cell is the activation of a cascade of primary and secondary target genes that ultimately drive the proliferation of infected cells. While infection of B cells does not require EBNA2, all subsequent steps including activation and cell cycle progression require the continuous presence of the EBNA2 protein. To study the impact of EBNA2-EBF1 complex formation on B cell immortalization, we established a recombinant EBV derivative with a deletion of the α1-helix in the END domain, termed EBVΔα1, and used it to infect primary human B cells. To examine B cell activation and the expansion of the cultures, MTT assays were performed on day 0 prior to infection and on day two, four, six, and eight post-infection (Figure 2A). On day two post-infection, MTT conversion of EBVwt and EBVΔα1 infected cell cultures was similar and modestly increased indicating that these B cells were activated compared to non-infected control cultures. Starting at day four post-infection, MTT conversion of EBVwt infected cells gradually increased reflecting the increase of total viable cells. In contrast, MTT conversion of EBVΔα1 infected B cells remained constant at low levels indicating that these cells remained viable for at least eight days but the cultures did not expand. To further characterize the proliferation defect of EBVΔα1 infected B cells, we analyzed the cell cycle phase distribution by bromodeoxyuridine (BrdU) incorporation and 7-AAD staining of total DNA followed flow cytometry (Figure 2B-D). Our gating strategy discriminated between G0/G1 (BrdU negative/DNA=2N), early S (BrdU medium/DNA=2N), advanced S phase (BrdU high/DNA≥2N) and G2/M phase (BrdU low/DNA≥2N) (Figure 2B). BrdU incorporation was detected as early as two days post-infection. EBVwt infected B cells progressed through the S phase and G2/M within four days and continued to cycle as expected (Mrozek-Gorska *et al*., 2019). In contrast, EBVΔα1 infected cells incorporated BrdU at low levels and maintained this state for at least eight days but cells were mostly arrested in G0/G1 and early S and barely advanced further (Figure 2C, D). Apoptotic cells accumulated in both EBVwt and EBVΔα1 infected cultures, but this was more pronounced in EBVΔα1 infected cultures.

**Figure 2.**
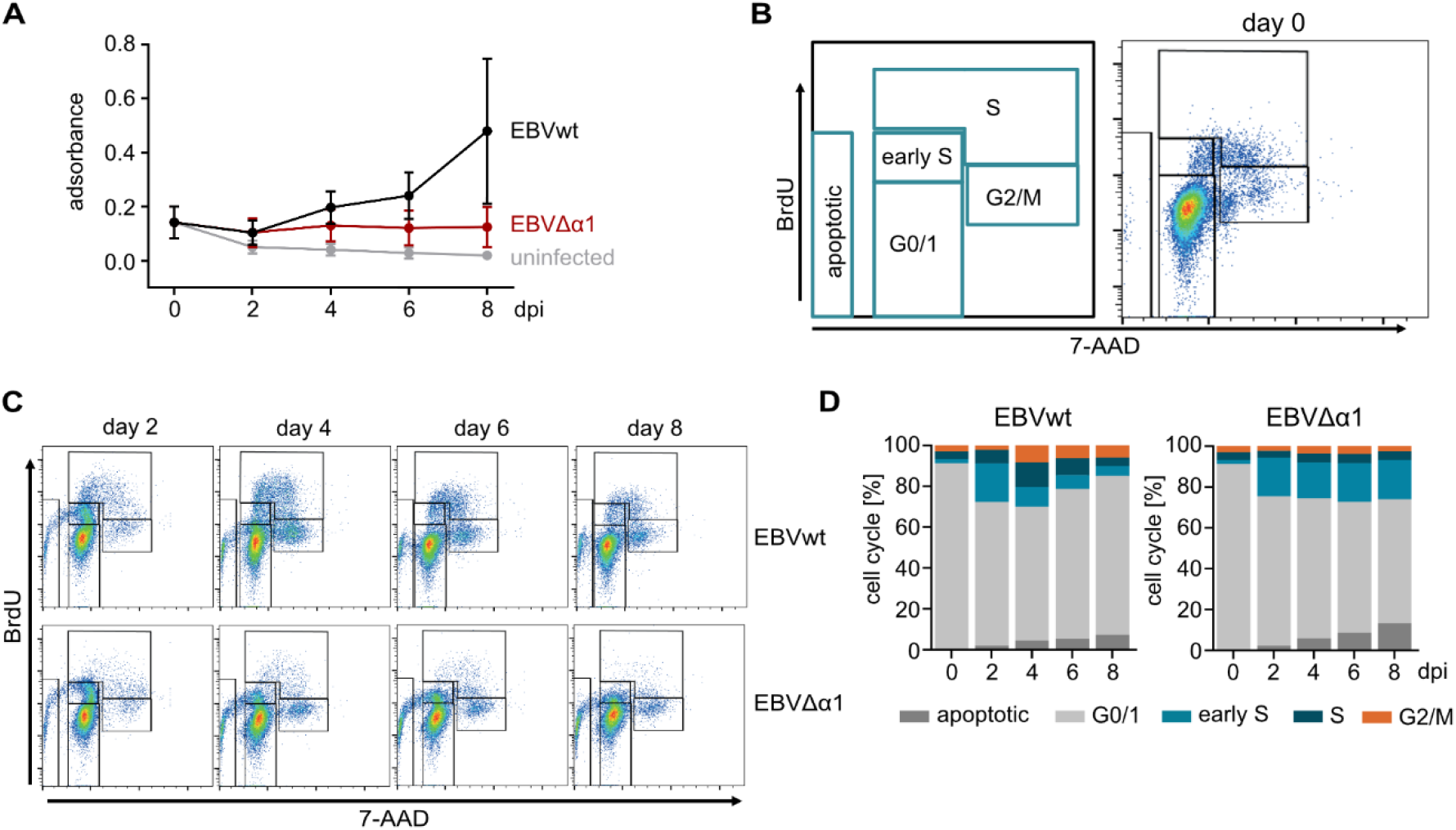
EBVΔα1 infected B cells arrest in the early S phase. (A) MTT assay of primary B cells infected with EBVwt or EBVΔα1 on day 0, two, four, six, eight post-infection. The mean of three biological replicates is plotted. Error bars indicate the standard deviation. Day 0 = non-infected B cells. (B) Gating strategy for the cell cycle analysis with bromodeoxyuridine (BrdU) and 7-AAD and one representative FACS plot for non-infected cells (day 0). All cells were gated on lymphocytes and single cells. (C) Results of the BrdU assays of one representative experiment. The assay was performed with B cells infected with EBVwt or EBVΔα1 on day two, four, six and eight post-infection. (D) Summary of the cell cycle analysis showing the mean results of three biological replicates. Day 0 = non-infected B cells; dpi – days post-infection.

### CD40 activation rescues long-term proliferation of EBVΔα1 infected B cells

Primary human B cells proliferate in response to CD40 activation in the presence of IL-4 (Banchereau *et al*., 1990). CD40 is a co-stimulatory receptor on the surface of B cells, which is triggered by CD40 ligands (CD40L) on helper T cells to activate canonical and non-canonical NFκB signaling, STAT5 phosphorylation and p38, AKT and JNK activation (Elgueta *et al*., 2009). Together, these signals resemble in many aspects the constitutive signaling of the viral LMP1 protein in EBV infected B cells and are critical for the survival of lymphoblastoid cell lines (LCL) evolving from EBV infected primary B cells *in vitro* (Thorley-Lawson, 2001). To study the phenotypes of long-term EBVΔα1 infected B cell cultures (LCLΔα1) and to establish them for further biochemical analyses, EBVwt and EBVΔα1 infected B cells were cultivated in the presence or absence of CD40L positive murine feeder cells in the absence of IL-4. In contrast to Banchereau et al, our model did not require IL-4 stimulation (Figure 3A) (Martina Wiesner *et al*., 2008). Non-infected B cells were used as control and cell cycle progression was analyzed by flow cytometry of propidium iodide (PI) stained nuclei.

**Figure 3.**
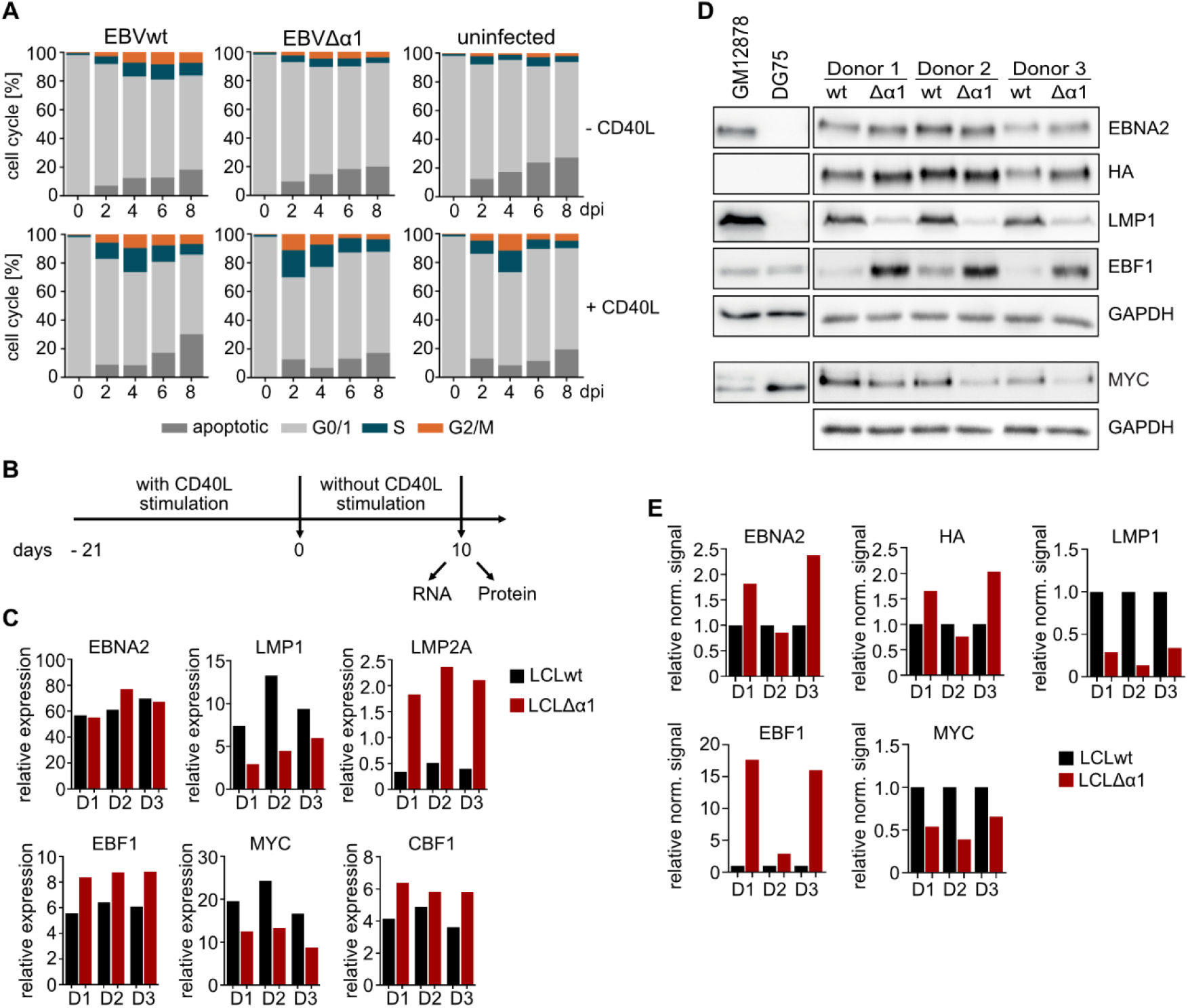
Long-term cultures of LCLΔα1 established by co-cultivation of primary B cells infected with EBVΔα1 on CD40 ligand (CD40L) feeder cells. (A) Cell cycle analyses of primary B cells infected with EBVwt or EBVΔα1. Non-infected or infected primary B cells were either cultured without (-) CD40L feeder cells or with (+) CD40L feeder cells. The cell cycle distribution was analyzed by propidium iodide staining (PI) staining and flow cytometry on day 0, two, four, six, eight post-infection. The mean of three biological replicates is plotted. See Figure S 2 for one representative experiment. (B) Experimental setup for RNA and protein preparation of LCLwt or LCLΔα1 cell lines established with the help of CD40L feeder cells were cultured without CD40L feeder cells for ten days prior to RNA and protein isolation. (C) RT-qPCR of viral and cellular genes in LCLwt or LCLΔα1 generated from three donors (D1, D2, D3) cultivated for 10 days without CD40L stimulus as in panel B. Transcript levels of viral and cellular genes were analyzed and normalized to RNA pol II transcript levels (see Figure S 3E for the mRNA expression level of RNA pol II) (D) Protein expression of latent viral and cellular proteins in LCLwt or LCLΔα1 generated from three donors (D1, D2, D3) after ten days without CD40L activation. Cell lysates of EBV infected GM12878 or EBV negative DG75 cells served as positive or negative controls, respectively. (E) Relative Western blot signals in (D) normalized to the corresponding GAPDH signals. (see Figure S 3F, G for details on GAPDH).

Similar to non-infected cells, EBVΔα1 infected B cells could not establish long-term cultures without CD40L feeder cells (Figure 3A). In contrast to feeder free cultures, all CD40L stimulated B cells entered the S and G2/M phases rapidly (Figure 3A). Remarkably, EBVΔα1 infected cells showed a dramatic increase in S phase already on day two post-infection and, overall, apoptosis was less pronounced in CD40L stimulated cells (Figure S 2). For comparative studies on RNA and protein expression, established pairs of LCLwt and LCLΔα1 derived from the same donor were cultivated in the absence of CD40L feeder cells for 10 days (Figure 3B). EBVΔα1 infected cells gradually ceased to proliferate but did not die, whereas proliferation of EBVwt infected cells was independent of CD40L stimulation as expected (Figure S 3A). While EBNA2 RNA abundance was similar in all three pairs of LCLwt and LCLΔα1, LMP1 was significantly decreased. Surprisingly, LMP2A was increased in LCLΔα1.

In these cells MYC levels were decreased, but EBF1 and CBF1 levels were increased (Figure 3C). The latent viral proteins EBNA2 and LMP1 were detected in all LCL. EBF1 protein expression was consistently elevated in all LCLΔα1 samples, while MYC and to an even stronger degree LMP1 were consistently reduced (Figure 3D, E). These data demonstrate that permanent CD40 ligation by feeder cells is required for the long-term survival of EBVΔα1 infected B cells.

It is known that both EBNA2 (Kaiser *et al*., 1999) and LMP1 (Dirmeier *et al*., 2005) induce *MYC*. We hypothesized that low LMP1 levels could be rate limiting for *MYC* expression causing the observed phenotypes in cell proliferation and survival of LCLΔα1 cells. We expressed LMP1 ectopically in EBVΔα1 infected cells but found that LMP1 did not restore their proliferation in the absence of CD40 stimulation (Figure S 3C, D). We concluded that elevated LMP1 expression levels are not sufficient to restore the proliferation of EBNA2Δα1 expressing cells.

### The α1-helix assists in EBF1 dependent EBNA2 chromatin binding

We have previously shown that EBNA2 can bind to chromatin in CBF1 knock-out cell lines (Glaser *et al*., 2017). This finding indicates that EBF1 and CBF1 may act independently to recruit EBNA2 to chromatin although the majority of EBNA2 binding sites encompass DNA sequence motifs that are bound by both EBF1 and CBF1. To study how the deletion of EBNA2’s α1-helix impairs the access of EBNA2 to chromatin, we performed chromatin immunoprecipitations followed by qPCR (ChIP-qPCR) on selected genomic regions using either EBF1 or EBNA2 specific antibodies. LCLs infected with EBVwt or EBVΔα1 were cultivated on CD40L feeder cells and were removed from feeder cell layers one day prior to chromatin isolation. The *HES1* gene, a well characterized canonical Notch target gene in T cells, carries two paired CBF1 binding sites within its promoter. They facilitate dimerization of intracellular Notch (Arnett *et al*., 2010) as Notch, similar to EBNA2, uses CBF1 as a DNA anchor (Strobl *et al*., 1997; Zimber-Strobl and Strobl, 2001). EBF1 is not expressed in T cells suggesting that HES1 activation by EBNA2 in B cells might be mediated solely by CBF1 and is independent of EBF1. Indeed, EBF1 did not bind to the HES1 promoter but EBNA2wt and EBNA2Δα1 did indicating that both proteins use CBF1 sites within the HES promoter to bind chromatin (Figure 4A, B). The experiments also indicated that the deletion of the α1 helix in EBNA2 does not affect CBF1 binding suggesting a correct folding of the mutant EBNA2 protein. CD2, a T cell marker gene, served as a negative control. As expected, neither EBF1 nor EBNA2 bound to this locus.

**Figure 4.**
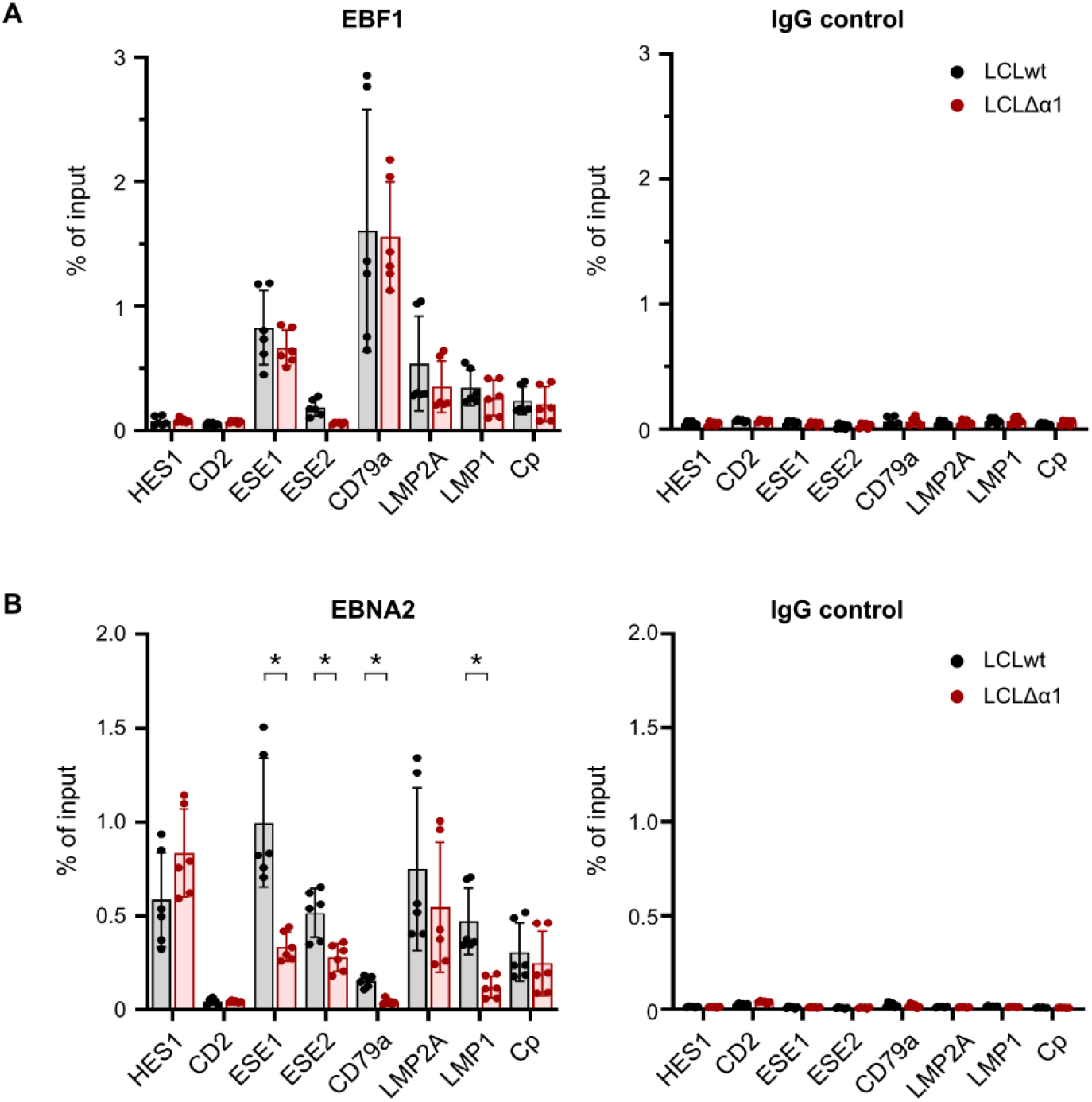
EBNA2’s α1-helix assists in EBF1 dependent EBNA2 chromatin binding. (A, B) ChIP-qPCR for (A) EBF1 or IgG control and (B) EBNA2 or IgG control in established LCLwt and LCLΔα1 at cellular and viral genomic regions. LCLwt and LCLΔα1 were cultured without CD40L expressing feeder cells for one day prior to chromatin preparation. CD2 was used as a negative control. Mean of three biological replicates is plotted with error bars indicating the standard deviation. Asterisks indicate p < 0.05. ESE – EBV super enhancer

The EBV super enhancers (ESE) 1 and 2 control *MYC* expression in EBV infected B cells (Zhou *et al*., 2015). EBF1 and EBNA2wt were recruited well to ESE1 but EBNA2Δα1 binding was significantly impaired indicating that EBNA2 uses EBF1 to interact with ESE1 (Figure 4A, B). EBNA2Δα1 recruitment to ESE2 was less pronounced compared to ESE1 but also significantly impaired compared to EBNA2wt. CD79a is an EBF1 target gene that is well characterized during B cell development (Hagman, Travis and Grosschedl, 1991). In LCLs, EBF1 bound strongly to the CD79a promoter. EBNA2 binding to the CD79a promoter was weak in comparison to other loci but strictly required the α1-helix within the END domain (Figure 4A, B). In addition, we tested the binding of EBNA2 at latent viral, EBNA2 responsive promoters. Activation of the LMP2A and the viral C-promoter (Cp) by EBNA2 is well characterized (Strobl *et al*., 1997) and strictly dependent on CBF1 (Zimber-Strobl *et al*., 1994) although EBF1 can be detected at these promoters. Both promoters, Cp and LMP2A, bound EBNA2Δα1 and EBNA2wt equally well and, thus, EBNA2 binding was independent of EBF1. These findings as well as elevated LMP2A expression levels in LCLΔα1 cells (Figure 3) are consistent with previous reports, that have described enhanced LMP2A expression in EBF1 depleted but diminished LMP2A expression in CBF1 depleted LCLs (Lu *et al*., 2016). Thus, despite elevated EBF1 levels in LCLΔα1 cells (Figure 3) and equal binding to the LMP2A promotor (Figure 4A), regulation of LMP2A was independent of EBF1. As reported previously (Zhao *et al*., 2011; Murata *et al*., 2016), EBF1 was recruited to the LMP1 promoter, but efficient EBNA2 binding was clearly impaired in the absence of the α1-helix (Figure 4B). In summary, EBNA2Δα1 is a highly specific loss-of-function mutant that selectively abolishes EBF1 dependent functions of EBNA2.

### EBVΔα1 fails to efficiently initiate the expression of critical cellular genes upon B cell infection

As illustrated in figure 2, EBVΔα1 infected B cells started to enter the S phase of the cell cycle on day two post-infection but progression through the S and G2/M phase in the following days was strongly impaired. To characterize the defect of this mutant in detail, we analyzed the gene expression patterns of naïve resting B cells prior to infection (day 0) and on day one, two, three, and four post-infection with either EBVwt or EBVΔα1. To this end, adenoid human B cells were isolated and sorted for naïve resting B cells (IgD+/CD38-) by flow cytometry (Figure 5A) (Mrozek-Gorska *et al*., 2019). Forward and side scatter of all cells indicated an increase in cell size and granularity on day two post-infection but these morphological changes were more pronounced in EBVwt infected B cells (Figure 5A).

**Figure 5.**
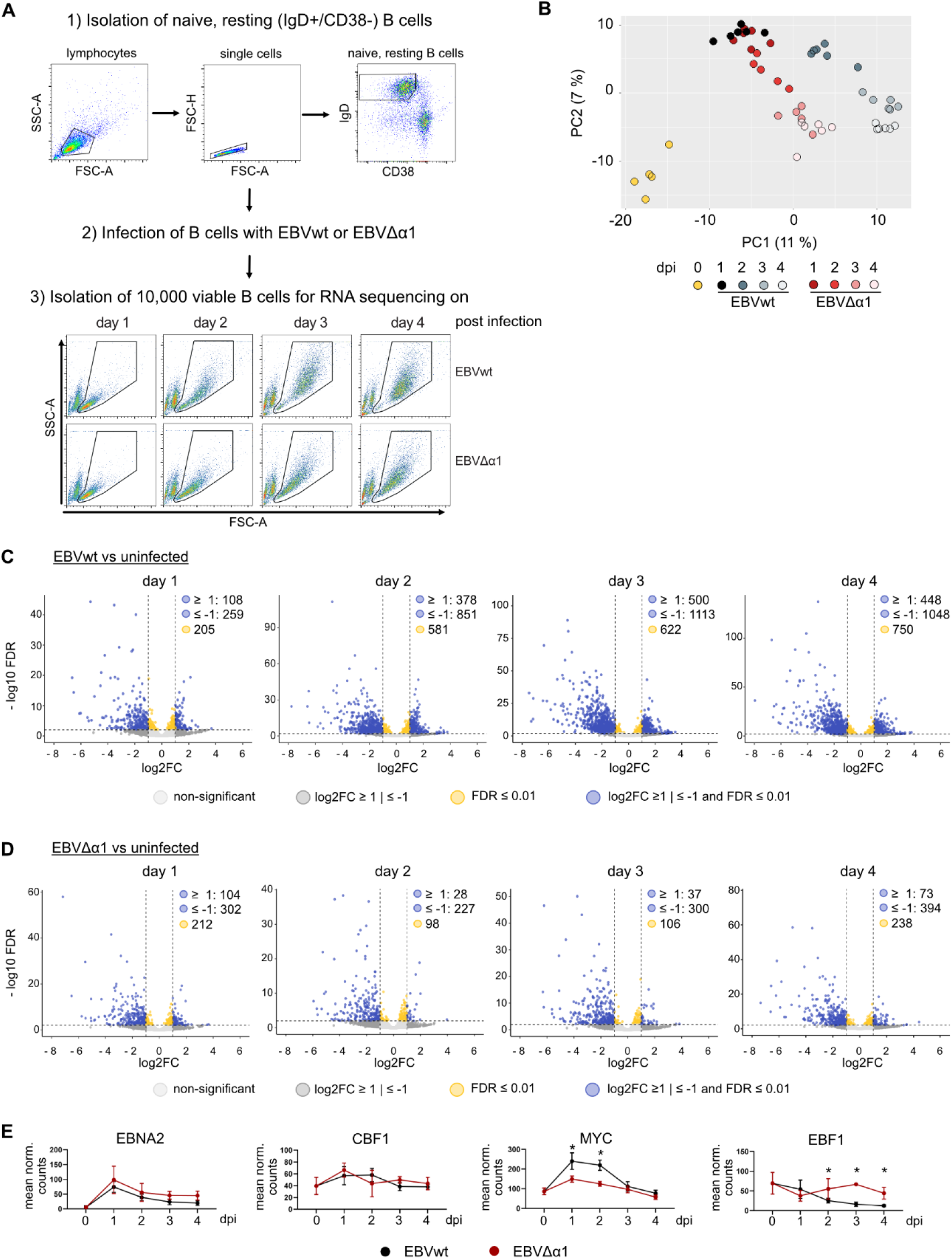
Dynamic changes of gene expression patterns in B cells infected with EBVwt or EBVΔα1 (A) Workflow for cell preparation and infection. Naïve resting B cells (IgD+/CD38-) were isolated from adenoids and infected with EBVwt or EBVΔα1. 10,000 non-infected, naïve resting B cells were collected to serve as the day 0 sample. On day one, two, three, four after infection, 10,000 viable cells were isolated by flow cytometry sorting and collected for RNA sequencing. Flow cytometry plots of one representative experiment are shown. (B) Principal component analysis of all protein coding genes. The percentage of variance explained by the first (PC1) and second (PC2) principal components are shown in parentheses. (C, D) Volcano plots of differentially expressed (DE) protein coding genes comparing (C) EBVwt or (D) EBVΔα1 infected B cells on day one, two, three, four post-infection to non-infected samples (day 0). Dotted lines indicate log2FC = 1 and FDR = 0.01. (E) Mean normalized expression of EBNA2, CBF1, MYC and EBF1 in EBVwt and EBVΔα1 infected B cells on day 0, one, two, three, four post-infection. Error bars indicate standard deviation and asterisks indicate FDR < 0.001 calculated by DESeq2.

Total cellular RNA was isolated on individual days and high quality cDNA libraries for RNA-sequencing analyses were generated (Figure S 4) using the bulk RNA-sequencing method prime-seq (Janjic *et al*., 2021). Principal component analysis (PCA) of RNA sequencing results for protein coding genes confirmed the considerable switch of gene expression patterns caused by viral infection of both viruses (Figure 5B). Moreover, the PCA illustrated a substantial overlap of the samples obtained with EBVwt and EBVΔα1 infection on day one and a stronger divergence starting from day two and thereafter (Figure 5B). Overall, the regulation of gene expression in B cells was affected to a greater degree upon EBVwt infection compared to EBVΔα1 infection on day two and thereafter (Figure 5C, D). Surprisingly, we detected a significant loss of transcripts over time in infected compared to non-infected cells which might have been caused by repression of specific genes or loss of transcript stability in activated cells (Figure 5C, D). Regarding viral genes, only 4920 reads could be mapped to the viral genome of which 3095 were mapped to EBNA2. This, unfortunately, provided us with little information about viral gene expression and EBNA2 was the only gene with meaningful expression levels. Expression of EBNA2 in EBVwt and EBVΔα1 infections was similar during the four days of the experiment (Figure 5E). Also, no significant differential expression of CBF1 was detected between viral strains over time (Figure 5E). MYC induction peaked in both infection conditions one day post-infection but was significantly less induced in EBVΔα1 infected cells (Figure 5E). In contrast, EBF1 transcript levels were significantly higher compared to EBVwt infected cells on day two, three, and four (Figure 5E). Taken together, the data show that gene expression was altered upon infection of B cells with EBVwt or EBVΔα1. However, the degree of gene regulation was strongly impaired in EBVΔα1 infected B cells although EBNA2 expression levels were comparable to EBVwt infected B cells.

### EBVΔα1 fails to initiate important processes required for cell cycle progression in infected B cells

We performed gene clustering with subsequent gene ontology (GO) term analysis on differentially expressed (DE) genes with an FDR < 0.1 and a fold change of ≥ 2 (Mrozek-Gorska *et al*., 2019) to study the dynamics of gene regulation and their functions. We could identify dynamic gene clusters with associated GO terms in EBVwt (Figure S 5B-F) and EBVΔα1 (Figure S 5G-J) infected cells. The gene cluster associated with the cell cycle and division had a similar dynamic (Figure S 5B, G), but it was larger in the EBVwt (1033 genes) than in EBVΔα1 (732 genes) infected cells. Another interesting observation was that genes associated with RNA metabolism (“wt dark blue” and “Δα1 light blue”, Figure S 5C, H) peaked on day one post-infection but tended to decrease more markedly in EBVΔα1 infected cells towards levels seen in non-infected cells (day 0). Importantly, the “wt dark red cluster”, strongly associated with biosynthesis and mitochondrial processes, did not show up in EBVΔα1 infected cells. In fact, most of the genes in this cluster were either not differentially expressed in the mutant (74.6 %) or they tended to be only transiently activated (i.e., 17.4 % of those genes are in the “Δα1 light blue” cluster, see Figure S 5H).

Next, we performed a gene set enrichment analysis (GSEA) with differentially expressed, protein coding genes (FDR < 0.01) to identify cellular processes affected in EBVΔα1 infected B cells at each time point. We included the molecular signature database hallmark gene sets for the GSEA, which encompassed two clusters for MYC regulated genes (MYC targets V1/V2) (Liberzon *et al*., 2015). MYC targets V1 defines a broader set of downstream targets whereas MYC targets V2 is more narrowly defined. Notably, several gene sets were exclusively enriched in EBVwt infections and included the MYC targets V2 as well as cellular and metabolic gene sets such as DNA repair, fatty acid metabolism, glycolysis or reactive oxygen species (ROS) pathways (Figure S 5K). GSEA with DE genes identified by comparing EBVΔα1 vs EBVwt infections revealed that further important processes required for a successful establishment of latency were more strongly enriched in EBVwt than the EBVΔα1 infected cells (Figure 6A). *MYC* is an important target gene of EBNA2 (Kaiser *et al*., 1999) and a stronger correlation of MYC targets V1/ V2 with EBVwt infection could indicate insufficient levels of MYC in EBVΔα1 infected cells. E2F targets and the G2M checkpoint include genes that are important for the cell cycle, which has been reported to be initiated between day three and four post-infection (Mrozek-Gorska *et al*., 2019) but we could detect an earlier cell cycle onset. Metabolic processes such as glycolysis, oxidative phosphorylation (OXPHOS), fatty acid metabolism, ROS pathways and mTORC1 signaling are pathways upregulated upon B cell activation (Akkaya and Pierce, 2019; Egawa and Bhattacharya, 2019). Gene sets significantly enriched in cells infected with either viruses on day one included MYC targets V1, E2F targets, OXPHOS, the G2M checkpoint and mTORC1 signaling (Figure 6B-K). Interestingly, the enrichment for MYC V1 was comparable in EBVwt or EBVΔα1 infected cells on day one but the normalized enrichment scores (NES) declined in EBVΔα1 infected B cells afterwards. The E2F targets, OXPHOS and G2M gene sets were only enriched in the EBVΔα1 infection at later time points indicating a delayed induction of genes from these sets. Additionally, all gene sets that were present in both infection conditions were more strongly enriched in the EBVwt infection than EBVΔα1 infection (Figure 6C, E, G, I). mTORC1 signaling was enriched in both, mutant and wild type infected cells on day one post-infection (Figure 6J, K). However, this important sensor for nutrients, redox state and energy supply was lost in EBVΔα1 infections in the following days. Thus, an important signal that activates mRNA transcription and protein translation was initiated in cells infected with both viruses but not maintained in cells infected with EBVΔα1. Overall, RNA sequencing revealed that changes in gene expression, especially starting on day two and onwards, were less pronounced in EBVΔα1 compared with EBVwt infected B cells and partially reminiscent of non-infected cells. Additionally, important target genes and metabolic pathways were not induced or delayed in EBVΔα1 infected cells.

**Figure 6.**
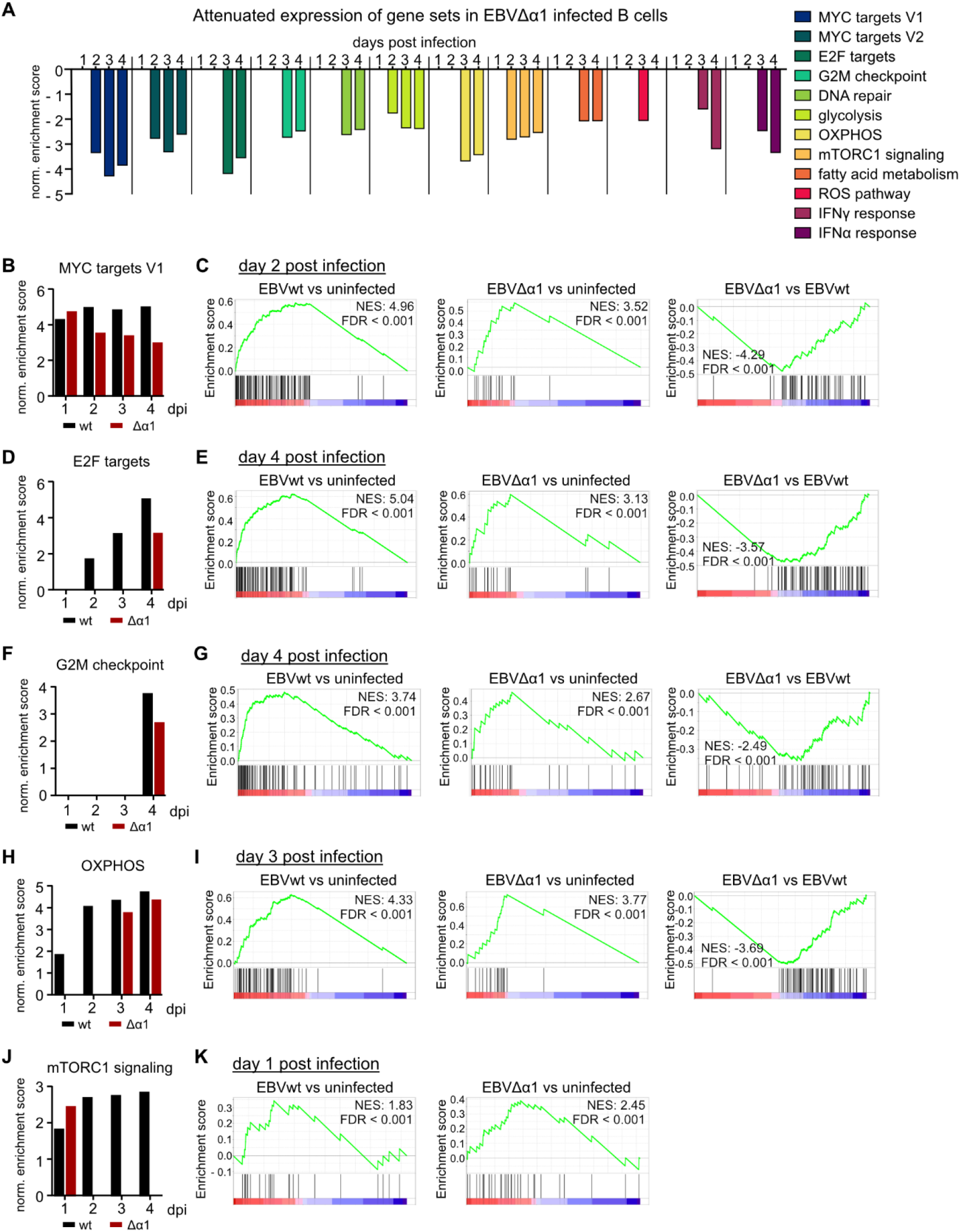
EBVΔα1 cannot efficiently induce critical cellular and metabolic processes. Gene set enrichment analysis (GSEA) was performed with differentially expressed (DE), protein coding genes that displayed a p-value of ≤ 0.01 as calculated by DESeq2. Gene sets with an FDR < 0.05 were considered significant. (A) GSEA with genes that were differentially expressed between EBVΔα1 and EBVwt infected B cells. The bar graph shows the NES for each gene set at each day post-infection. A negative NES correlates with an enrichment in EBVwt infected B cells over EBVΔα1 infected B cells. (C-I) Gene sets that are present in both EBVwt and EBVΔα1 infected B cells at a least one day post-infection. Bar graphs are showing the NES at each day post-infection for the gene sets (B) MYC targets V1 (D) E2F targets (F) oxidative phosphorylation (OXPHOS) (H) G2M check point and (J) mTORC1 signaling. DE genes of the analysis EBVwt vs non-infected or EBVΔα1 vs non-infected were included for these GSEA. Enrichment plots of the GSEA with DE genes for the indicated analysis display (C) MYC targets V1 on day two post-infection (E) E2F targets on day four post-infection (G) OXPHOS on day 3 post-infection (H) G2M checkpoint on day four post-infection and (K) mTORC1 signaling on day one post-infection.

## DISCUSSION

### EBVΔα1 can activate primary human B cells but cell cycle progression is aborted in the early S phase

Upon infection of primary resting B cells, EBV establishes a complex specific cellular and viral gene expression program. Eventually, a limited number of viral latent nuclear antigens (EBNAs) and latent membrane proteins (LMPs) cooperate to establish the long-term proliferation of infected B cell cultures. EBV devoid of EBNA2 (Pich *et al*., 2019) or EBV mutants with EBNA2 alleles incapable of interacting with CBF1 (Yalamanchili *et al*., 1994) cannot activate infected B cells demonstrating the importance of EBNA2 and its association with CBF1 for the immortalization process. EBNA2 is a multifunctional viral oncogene that is associated with a plethora of cellular proteins (Kempkes and Ling, 2015). While the core of the EBNA2 binding site of CBF1 has been carefully mapped to a central double tryptophan motif (Ling, Rawlins and Hayward, 1993), information on the exact binding motif of further EBNA2 binding partners is limited and recombinant viral mutants, which lack specific binding sites, have not been reported.

We have shown previously that EBNA2 requires its N-terminal dimerization domain (END) to build stable complexes with the early B cell factor (EBF) 1, a candidate factor to confer B cell specificity to EBNA2 (Glaser *et al*., 2017). Here, we characterize an EBNA2 mutant (EBNA2Δα1) lacking the α1-helix, which consists of 5 amino acids, within the EBNA2 N-terminal dimerization (END) domain and show that EBNA2Δα1 is deficient for EBF1 binding. To study the impact of the association of EBF1 with EBNA2 on B cell immortalization, we generated viral mutants and found that a mutant EBV encoding EBNA2Δα1 (EBVΔα1) failed to immortalize primary human B cells. B cells infected with EBVΔα1 became activated and entered the S phase but failed to complete the cell cycle (Figure 2) causing a growth arrest that was rescued by CD40 stimulation (Figure 3A). The phenotype of EBVΔα1 infected B cells differs dramatically from B cells infected with an EBNA2 knockout EBV. B cells infected with EBNA2 knockout EBV do not survive until three days post-infection and fail to synthesize cellular DNA (Pich *et al*., 2019).

### EBNA2Δα1 selectively impairs EBF1 related functions of EBNA2

In EBV infected B cells, EBNA2 directly drives the expression of LMP1 (Grossman *et al*., 1994; Laux *et al*., 1994; Waltzer *et al*., 1994; Johannsen *et al*., 1995; Kaiser *et al*., 1999). In LCLs, roughly 50% of LMP1 promoter activation by EBNA2 depends on CBF1 (Yalamanchili *et al*., 1994). Multiple studies (Zhao *et al*., 2011; Lu *et al*., 2016; Murata *et al*., 2016) have demonstrated that EBF1 is critical for EBNA2 dependent LMP1 activation. Using the EBF1 binding deficient EBVΔα1 mutant virus, we demonstrate that the activation of LMP1 by EBNA2 requires the interaction with EBF1 via the α1-helix of the END domain. Recruitment of EBNA2Δα1 to the LMP1 promotor was reduced (Figure 4) leading to reduced LMP1 expression level in cells infected with the EBNA2Δα1 mutant EBV (Figure 3). In contrast, EBNA2Δα1 bound equally well to the CBF1 dependent viral C-, LMP2A, and cellular HES1 promoter as EBNA2wt did. Thus, EBVΔα1 carries a specific loss-of-function mutation in EBNA2 that does not affect functions related to the CBF1 interaction but selectively affects EBNA2’s transactivation capacity linked to EBF1 (Figure 4).

### Expression of MYC and basic cellular processes are impaired in EBVΔα1 infected B cells

EBNA2 is involved in all events of B cell activation and survival *ex vivo* because EBNA2 directly activates MYC expression (Kaiser *et al*., 1999). The central role of MYC in LCLs is well documented as ectopic expression of *MYC* can drive the proliferation of latently infected B cells even when EBNA2 is inactivated (Polack *et al*., 1996). EBNA2 regulates *MYC* by inducing chromatin loops that bridge two enhancers located – 556kb (ESE1) and – 428kb (ESE2) upstream of the *MYC* transcriptional start site to the *MYC* promotor (Zhao *et al*., 2011; McClellan *et al*., 2013; Wood *et al*., 2016; Jiang *et al*., 2017) EBF1 is recruited to both enhancers, although ESE1 is more active than ESE2 (Zhou *et al*., 2015). We confirm this differential binding pattern of EBF1 in our study (Figure 4). Importantly, EBNA2Δα1 binding was significantly impaired at ESE1 and ESE2 demonstrating that the α1-helix, consisting of 5 amino acid residues, substantially contributes to EBNA2 binding to both B cell specific *MYC* enhancers. This nicely explains significantly reduced MYC levels one day post-infection and reduced MYC levels in established LCLΔα1.

In LCLs, LMP1 enhances *MYC* expression in the presence of EBNA2 (Dirmeier *et al*., 2005) and thereby contributes to the proliferation of LCLs. We tested if ectopic expression of LMP1 in long-term LCLΔα1 cultures (Figure 3, Figure S 3) could substitute for CD40 stimulation (Figure S 3) but found that ectopic LMP1 expression was not sufficient to promote the proliferation of LCLΔα1 cultures. We conclude, that cellular pathways downstream of CD40 signaling, which are not provided by LMP1, are important to maintain continuous proliferation of these cultures. Since we know that high level MYC will certainly rescue any EBNA2 deficiency in LCLs (Polack *et al*., 1996), we did not test if ectopic expression of *MYC* might rescue the proliferation of LCLΔα1. Such experiments would not provide new mechanistical insights into the specific function of EBNA2-EBF1 complexes. In summary, we conclude that full *MYC* activation by EBNA2 is achieved through the interaction with EBF1 allowing EBNA2 the access to specific *MYC* enhancers early after infection of naïve B cells as well as in established LCLs. Since EBNA2 (Wu *et al*., 1996; Wood *et al*., 2016) and EBF1 (Gao *et al*., 2009; Wang *et al*., 2020) interact with the chromatin remodeler Brg1, we speculate that this interaction might contribute to the regulation of the EBV specific MYC enhancers and, thereby, might enhance MYC activation.

To characterize the defects of EBVΔα1 infected B cells in detail, a time course RNA-sequencing experiment was performed surveying non-infected cells and the subsequent early phase until day four post-infection. Similar patterns of gene expression changes were grouped into clusters for EBNA2wt and EBNA2Δα1 expressing cells and a gene ontology term analysis was performed for each cluster (Figure S 5B-J). Cell cycle and RNA processing functions as well as multiple biosynthetic pathways were attenuated in EBVΔα1 infected cells. According to the gene set enrichment analysis, several enriched gene sets were exclusively found in EBVwt but not in EBVΔα1 infected B cells (Figure S 5K). Induction of MYC and E2F target genes, both critical for G1 phase cell cycle progression, was severely impaired in EBVΔα1 infected B cells already one day post-infection. Also, glycolysis, oxidative phosphorylation, and fatty acid metabolism, which are known to be closely linked to distinct phases of the cell cycle (Icard *et al*., 2019), were attenuated in these cells. In summary, the phenotype of EBVΔα1 infected B cells revealed a broad spectrum of alterations in gene expression that can be well explained by low levels of MYC in these cells. MYC is a pleiotropic transcription factor that not only directly activates target gene expression by binding to specific DNA sequences of regulatory genomic sites but, in addition, accumulates at active promoter sites and amplifies the transcriptional activity of a global spectrum of target genes (Nie *et al*., 2012). In cancer cells, oncogenic MYC levels are implicated in many metabolic pathways to regulate and fulfill the increased demand for energy and compounds for cell growth and proliferation (Dong *et al*., 2020). Since hyperactivation of *MYC* is absent in EBVΔα1 infected B cells (Figure 5E), the metabolic defects in these cells are most likely caused by the inefficient activation of *MYC*.

During development, EBF1 acts as a pioneer factor that alters the chromatin signature of lymphocyte precursors directing them into the B cell linage (Boller *et al*., 2016). In murine pro-B cells EBF1 activates *MYC* and promotes proliferation of these cells (Somasundaram *et al*., 2021). Our study highlights how EBF1, in cooperation with EBNA2, supports the activation of basic metabolic processes in human peripheral mature B cells. In conclusion, the protruding α1-helix on the surface of the EBNA2 N-terminal dimerization (END) domain is a key structural element of EBNA2 that confers B cell proliferation as a first step towards establishment of latency by high-jacking developmental signaling cues.

## Supporting information

Supplemental table 6 - cluster analysis EBVmt infection

Supplemental table 7 - cluster analysis EBVwt infection

Supplemental table 8 - normalized expression data of protein coding genes

## Author contributions

Designed research: SB, BK

Performed research: SB, LEW, XZ, WH, CKR

Analyzed data: SB, AS, LEW, XZ, CKR, WH, WE, BK

Wrote the manuscript: SB, AS, BK

## Acknowledgement

We thank Ezgi Akidil, Dagmar Pich and Adam Yen-Fu Chen (HMGU) for valuable experimental support, LAFUGA (Ludwig Maximilians University) for the excellent sequencing service and the monoclonal antibody core facility (HMGU) for providing antibodies.

LEW and WE were supported by the Deutsche Forschungsgemeinschaft (DFG) through the SFB1243 (Subproject A14) and the Cyliax foundation. XZ was supported by the China Scholarship Council (CSC No.: 201603250052).

## MATERIAL and METHODS

### MATERIALS

**Table 1.**
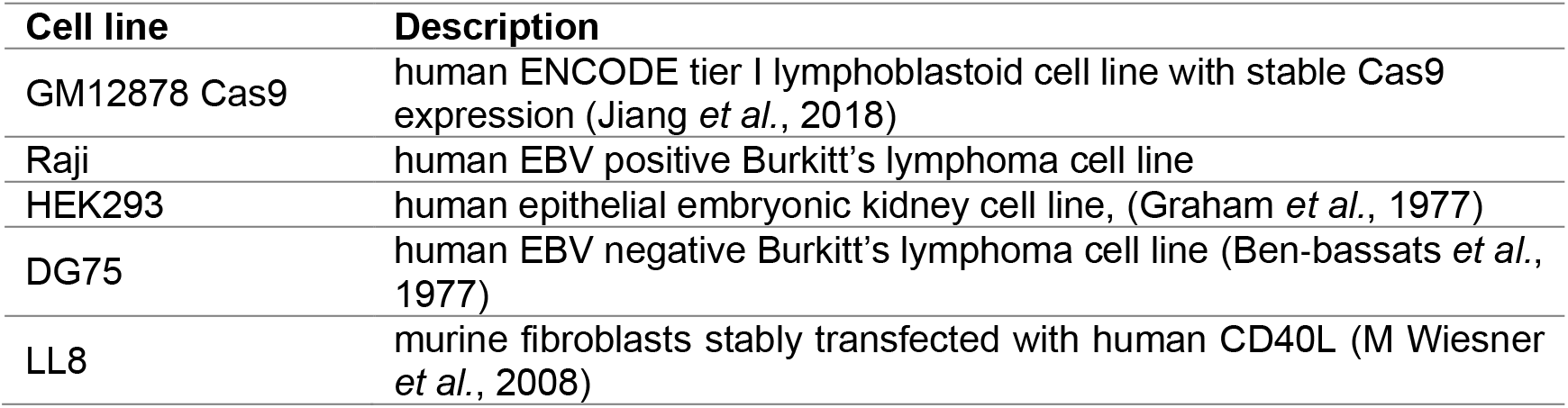
General cell lines

| Cell line | Description |
| --- | --- |
| GM12878 Cas9 | human ENCODE tier I lymphoblastoid cell line with stable Cas9 expression (Jiang *et al.*, 2018) |
| Raji | human EBV positive Burkitt's lymphoma cell line |
| HEK293 | human epithelial embryonic kidney cell line, (Graham *et al.*, 1977) |
| DG75 | human EBV negative Burkitt's lymphoma cell line (Ben-bassats *et al.*, 1977) |
| LL8 | murine fibroblasts stably transfected with human CD40L (M Wiesner *et al.*, 2008) |

**Table 2.** Lymphoblastoid cells lines established from donors

| Cell line | Donor | Virus |
| --- | --- | --- |
| SB268 LCLwt | 1 | XZ143 EBV (6008) HA-EBNA2 |
| SB301 LCL$\Delta\alpha 1$ | | SB161 EBV (6008) HA-EBNA2$\Delta\alpha 1$ |
| SB346.1 LCLwt | 6 | XZ143 EBV (6008) HA-EBNA2 |
| SB346.2 LCL$\Delta\alpha 1$ | | SB161 EBV (6008) HA-EBNA2$\Delta\alpha 1$ |
| CKR654.1 LCLwt | 8 | XZ143 EBV (6008) HA-EBNA2 |
| CKR654.3 LCL$\Delta\alpha 1$ | | SB161 EBV (6008) HA-EBNA2$\Delta\alpha 1$ |

**Table 3.** Conditional cell lines

| Cell line | Internal designation | Description |
| --- | --- | --- |
| DG75 doxHA-EBNA2 | CKR128-34 | human EBV negative Burkitt's lymphoma cell line carrying the doxycycline inducible pRTR-HA-EBNA2 plasmid |
| DG75 doxHA-EBNA2ΔEND | CKR505.11 | human EBV negative Burkitt's lymphoma cell line carrying the doxycycline inducible pRTR-HA-EBNA2ΔEND plasmid |
| DG75 doxHA-EBNA2-H15A | CKR606.17 | human EBV negative Burkitt's lymphoma cell line carrying the doxycycline inducible pRTR-HA-EBNA2-H15A plasmid |
| DG75 doxHA-EBNA2Δα1 | CKR607.20 | human EBV negative Burkitt's lymphoma cell line carrying the doxycycline inducible pRTR-HA-EBNA2Δα1 plasmid |
| LCLwt_doxLMP1 | CKR700.1 | LCL generated from adenoid biopsy, infected with EBVwt, stably transfected with pRTS-HA-LMP1 |
| LCLΔα1_doxLMP1 | CKR724.1 | LCL generated from adenoid biopsy, infected with EBVΔα1, stably transfected with pRTS-HA-LMP1 |

**Table 4.**
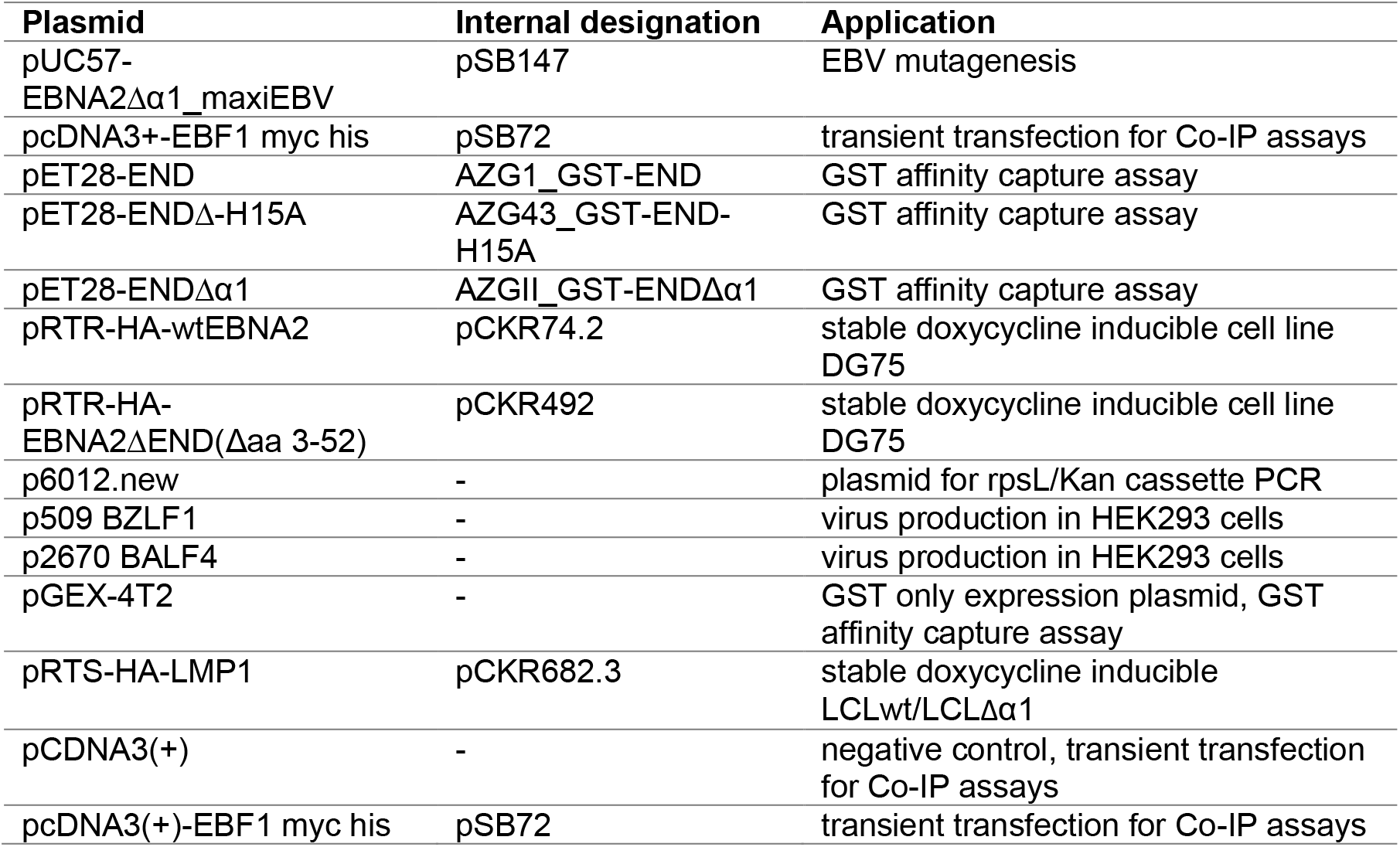
Plasmids

**Table 5.**
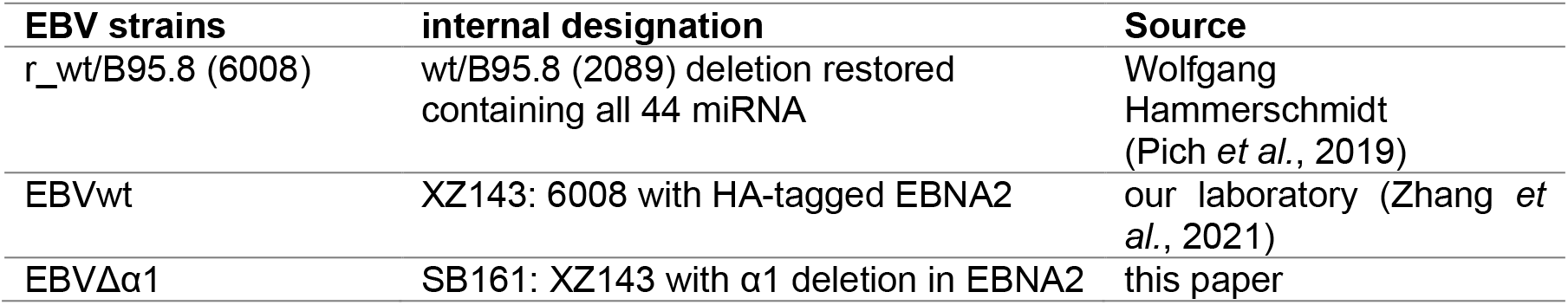
EBV strains

**Table 6.**
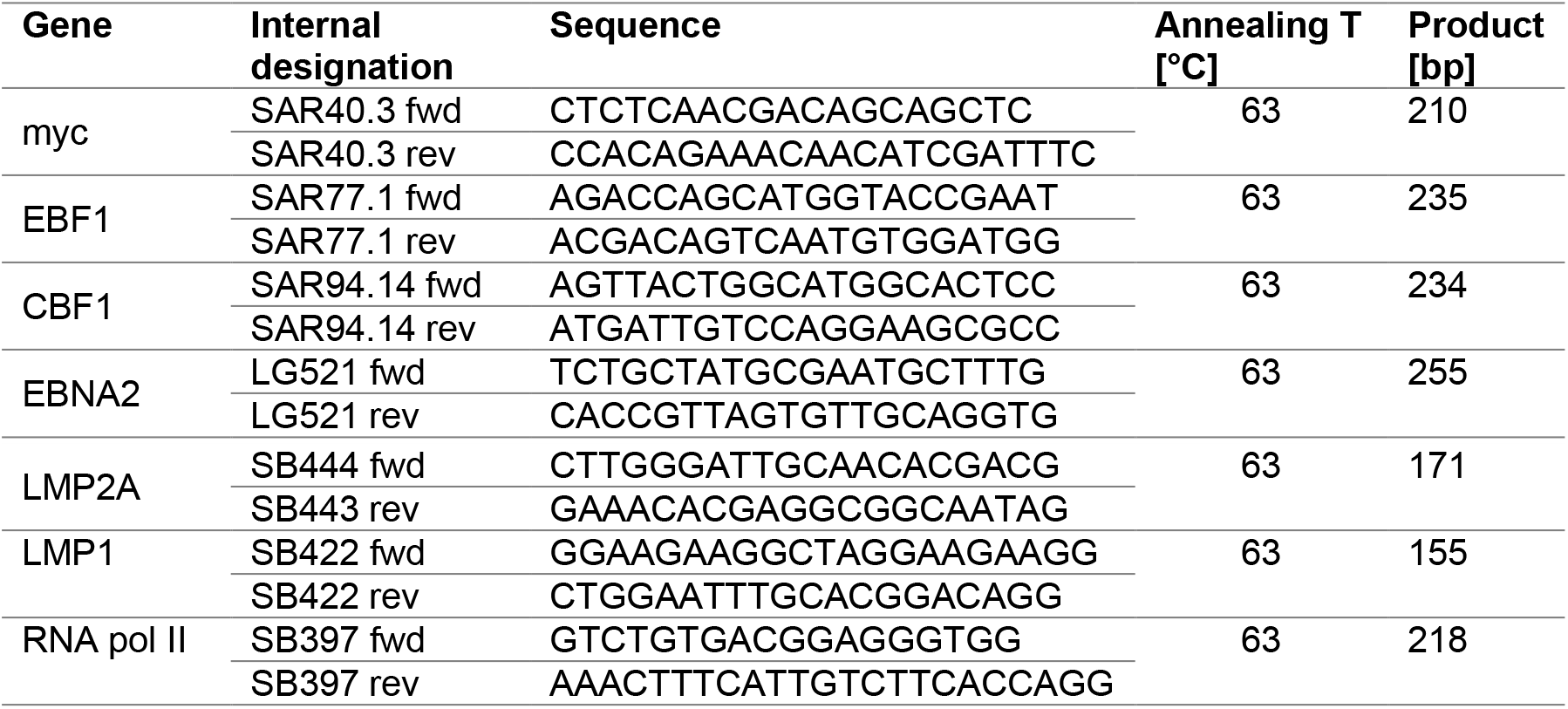
qPCR primer for cDNA quantification

**Table 7.**
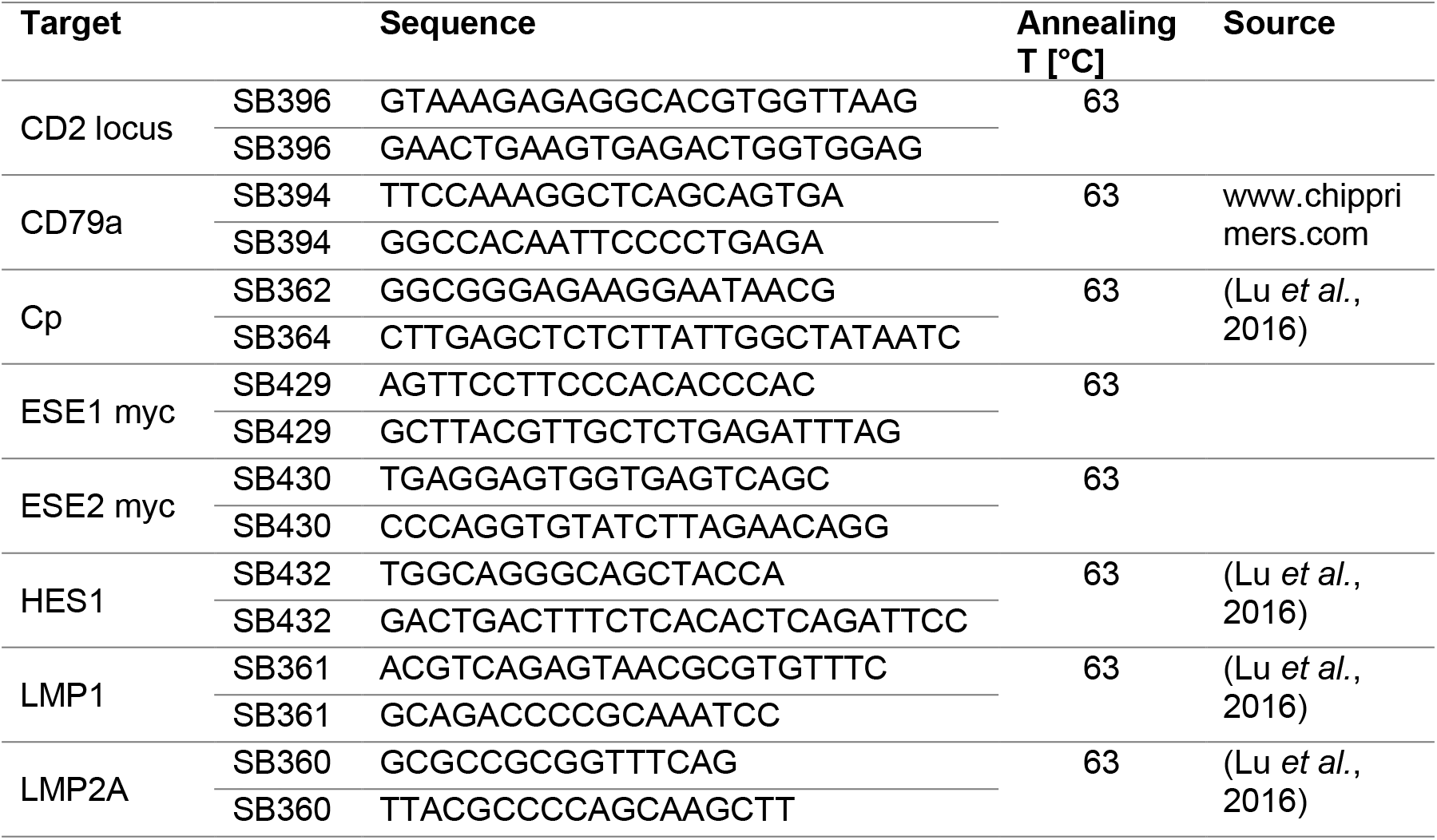
qPCR primers for ChIP quantification

**Table 8.** Critical commercial assays

| Product | Source | Identifier |
| --- | --- | --- |
| BrdU-APC Flow Kit | BD Pharmingen | cat. no. 552598 |
| ECL | GE Healthcare (Amersham) | cat. no. RPN2109 |
| NucleoSpin™ Gel and PCR Clean-up | Macherey & Nagel | cat. no. 740609.50 |
| LightCycler 480 SYBR Green I Master | Roche Diagnostics | cat. no. 04 887 352 001 |
| QIAshredder | Qiagen, Germany | cat. no. 79654 |
| RNase-free DNase Set | Qiagen, Germany | cat. no. 79254 |
| RNeasy Mini Kit | Qiagen, Germany | cat. no. 74104 |
| SuperScript IV First-Strand cDNA Synthesis Reaction | ThermoFischer | cat. no. 18091150 |
| Phusion®High-Fidelity Polymerase | New England Biolabs | cat. no. M0530 |
| NucleoBond™ Xtra BAC Maxi kit | Macherey-Nagel | cat. no. 740414 |

**Table 9.** Antibodies

| Specificity | Host | Application | Supplier |
| --- | --- | --- | --- |
| α-EBF1 (C-8) | mouse, IgG2a | WB, ChIP | Santa Cruz, cat# sc-137065 |
| α-EBF1 | rabbit, | ChIP | Sigma Aldrich, cat# ABE1294 |
| α-EBNA2 (R3) | rat, IgG2a | WB, IP, ChIP | HMGU Monoclonal Antibody Core Facility |
| α-EBNA2 (1E6) | rat, IgG2a | ChIP | HMGU Monoclonal Antibody Core Facility |
| α-EBNA3A (4A5-1111) | rat, | WB | HMGU Monoclonal Antibody Core Facility |
| α-EBNA3B (AN 6C9-1-1) | rat, | WB | HMGU Monoclonal Antibody Core Facility |
| α-EBNA3C (A10 P2-583) | rat, | WB | HMGU Monoclonal Antibody Core Facility |
| α- GST (6G9) | rat, IgG2a | WB, affinity capture, ChIP isotype control | HMGU Monoclonal Antibody Core Facility |
| α-Dog CD3 | rat, IgG1 | ChIP | HMGU Monoclonal Antibody Core Facility |
| α-HA (3F10) | rat, IgG1 | WB, ChIP | HMGU Monoclonal Antibody Core Facility |
| α-GAPDH (0411) | mouse, IgG1 | WB | Santa Cruz, cat# sc-47724 |
| α-LMP1 (1G6) | rat, | WB | HMGU Monoclonal Antibody Core Facility |
| α-myc (9E10) | mouse, | WB | HMGU Monoclonal Antibody Core Facility |
| α-IgD-FITC | mouse, IgG2a | FACS sorting | BD Biosciences, USA |
| α-CD38-PE-Cy7 | mose, IgG1κ | FACS sorting | eBioscience, cat #25-0389-42 |
| rabbit IgG |  | ChIP | Merck cat# PP64 |

### METHODS

#### Cell lines and routine cell culture conditions

All cells were cultured at 37 °C, 6 % CO_2_. DG75 (Ben-bassats *et al*., 1977) and irradiated LL8 (Martina Wiesner *et al*., 2008) cells were cultured in RMPI 1640 supplemented with 10 % FCS, 4 mM L-Glutamine, 100 U/ml penicillin, 100 μg/ml streptomycin. LCLwt_doxLMP1 and LCLΔα1_doxLMP1 were cultured with additional 40 μg/ml hygromycin.

Raji (Pulvertaft, 1964), HEK293 (Graham *et al*., 1977), primary B cells infected with EBVwt or EBVΔα1 and lymphoblastoid cell lines generated from EBVwt (LCLwt) or EBVΔα1 infection (LCLΔα1) were cultured in RPMI 1640 medium supplemented with 10 % FCS, 100 U/ml penicillin, 100 μg/ml streptomycin, 1 mM sodium pyruvate, 100 nM sodium selenite, and 0.43 %α-thioglycerols in 20 μM BCS. Primary B cells and LCLΔα1 were cultured on LL8 cells in LCL specific medium.

#### Irradiation of CD40L expressing LL8 feeder cells

A. Moosmann (Helmholtz Center Munich) provided irradiated CD40L expressing LL8 feeder cells. In brief, trypsinized cells were washed and resuspended in medium and irradiated with 180 Gy (γ-rays). 10^6^ cells were seeded per one full plate (96, 48, 24, 12, 6-well). 1.5×10^6^ cells were seeded for a T75 cell culture flask. LL8 cells were ready to use the next day.

#### Isolation of B cells from human adenoid tissue

To isolate primary B cells, adenoids were rinsed in medium and carefully pushed through a 100 μm cell strainer using a 10 ml syringe plunger. Depending on the adenoid size, the cell suspension was filled up to 25 ml or 50 ml with medium (RPMI 1640, 10% FCS, 100 U/ml penicillin, 100μg/ml streptomycin, 4 mM L-Glutamine). T cell rosettes were formed by adding 0.5 ml of sheep blood per 25 ml of cell suspension. 25 ml of the cell suspension with sheep blood was carefully layered onto 20 ml of Ficoll and centrifuged at 666 g, 40 min, at 10 °C, decelerated without breaks. The interface was transferred to a new 50 ml Falcon and washed three times with 40 - 50 ml PBS/2 mM EDTA at (i) 666 g (ii) 527 g (iii) 403 g, 10 min each, at 10 °C. Erythrocytes were lysed by resuspending the cell pellet in 5 ml red blood cell lysis buffer (154 mM NH_4_Cl; 9.98 mM KHCO_3_; 0.127 mM ETDA, pH8) followed by centrifugation for five minutes at 403 g, 10 °C. Cells were resuspended in LCL specific medium.

#### Primary B cell infection and generation of LCL

10^6^ primary B cells were infected with an MOI 0.1 with EBVwt or EBVΔα1 supernatant and cultured in LCL medium. For the first one to two weeks, 0.5 μg/ml Cyclosporin A and 10 μg/ml Ciprobay was added. To enhance the growth after one week of infection, B cells infected with EBVΔα1 were constantly cultured on irradiated LL8 cells. B cells infected with wild type EBV were kept without LL8 cells. Infected B cells were expanded to generate LCLs and frozen for long-term storage. Infected B cells for RNA sequencing were not cultured with LL8 cells.

#### Sorting of B cells to obtain subpopulations

10^8^ isolated, primary B cells were washed once with FACS buffer (PBS, 2% FCS, 2 mM EDTA) and resuspended in 1 ml FACS buffer. The cells were stained with 80 μl α-IgD-FITC (BD Pharmingen, #555778) and 24 μl α-CD38-PE-Cy7 (eBioscience, #25-0389-42) for one hour in the dark at 4 °C and washed twice with FACS buffer. The stained cells were resuspended in 6 ml FACS buffer and filtered through a 35 μm filter to obtain a single cell suspension. Sorting was performed on a FACS Aria IIIu device using the 70 μm nozzle, the “Purity” sorting mask and a sorting velocity of around 7000 events/second. The gating strategy included (i) lymphocytes (ii) single cells (iii) IgD+ CD38-. Sorted naïve resting B cells were used for infection experiments. To sort viable cells post-infection, the infected B cells were washed, resuspended in FACS buffer and filtered through a 35 μm strainer. Sorting was performed using a 100 μm nozzle, a velocity of around 7000 events/second with the “Purity” sorting mask. The gating was based on the forward scatter (FSC) and side scatter (SSC).

#### Cell cycle analysis with BrdU

Isolated, primary B cells were infected with an MOI 0.1 with EBVwt or EBVΔα1 and seeded with a density of 10^6^ cells/ml to analyze them on day 0, two, four, six and eight post-infection. Prior to cell harvest, the infected cells were pulsed with a final concentration of 10 μM BrdU for one hour at 37 °C. Afterwards, the cells were harvested, washed with 1 ml of FACS staining buffer (PBS, 2 % FCS, 2 mM EDTA) and centrifuged at 500 g for 5 min. The cells were fixed and permeabilized with 100 μl of BD Cytofix/Cytoperm for 25 min at RT and washed with 1 ml FACS buffer. The samples were stored in freezing medium (90% FCS, 10 % DMSO) at – 80°C until samples from all time points were collected. On the day of FACS analysis, freshly thawed samples were washed in FACS buffer, re-fixed with 100 μl BD Cytofix/Cytoperm for 10 min at RT and washed with 1 ml 1X BD Wash/Perm Buffer. Cell pellets were resuspended in 100 μl DNase (300 μg/ml in PBS final concentration) and incubated for one hour at 37 °C, 6% CO_2_. Afterwards, the cells were washed in 1 ml 1X BD Perm/Wash Buffer. Incorporated BrdU was stained with 1 μl α-BrdU-APC in 50 μl BD Perm/Wash buffer for 20 min in the dark at RT. After washing with 1 ml 1X BD Perm/Wash buffer, cells were resuspended in 20 μl 7-AAD solution, incubated for five minutes and filled up to 200 to 300 μl with FACS staining buffer. The analysis was done with FACS Fortessa.

#### Cell cycle analysis with PI

Isolated, primary B cells were infected with an MOI 0.1 with wild type EBV or EBVΔα1 and seeded with a density of 10^6^ cells/ml on plates either containing LL8 feeder cells or on plates without LL8 feeder cells. Cells were harvested on day 0, two, four, six and eight post-infection in preparation of the PI staining. 10^6^ cells were harvested and washed once in staining buffer (PBS, 2 % FCS). The cell pellet was fixed by adding 1 ml of ice-cold 70 % ethanol dropwise and stored at −20 °C until all samples from all time points were collected. On the day of cell cycle analysis, the fixed cell pellets were washed once in staining buffer. The cells were resuspended in 500 μl PI solution (10 μg/ml PI, 175 μg/ml RNaseA in PBS) and analyzed by flow cytometry.

#### Cell viability analysis with MTT

Isolated, primary B cells were infected with an MOI 0.1 with EBVwt or EBVΔα1. Non-infected B cells were included as a negative control. For each time point and condition, 8 replicates were prepared on a 96-well plate. 10^5^ cells/well were seeded in 100 μl medium. The MTT assays were performed on day 0, two, four, six and eight post-infection. 10 μl of MTT (5 mg/ml in PBS) was added to each well and incubated for four hours at 37 °C. Plates with MTT were stored at – 20 °C until measurement. On the day of analysis, formazan crystals were dissolved with 200 μl/well of 1 N HCl in isopropanol and the absorption was measured at 550 nm (measurement wavelength) and 690 nm (reference wave length). The background signal from medium without cells with MTT was subtracted from the sample values.

#### Virus production

In brief, HEK293 producer cells were seeded on dishes with a density of around 40 % one day before transfection. The medium was replaced with 25 ml puromycin-free medium just before the transfection. Component A (12 μg 1:1 BZLF1:BLAF4 expression plasmids, 1.2 ml RPMI 1640) and component B (72 μl PEI (1 mg/ml), 1.2 ml RPMI 1640) of the transfection solution were mixed and incubated for 15 min at RT. The solution was added dropwise to the cells and incubated for 3 days. The supernatant was harvested, centrifuged at 1200 rpm for 10 min and filtered through a sterile 1.2 μm filter. The virus titer of the supernatant was measured using Raji cells.

#### Virus titer calculation

Since the recombinant virus encodes for a GFP protein, Raji cell infections were used to calculate the titer of all virus supernatants. 10^5^ Raji cells were infected with 1, 2, 5, 10 and 20 μl in a 24-well plate and incubated for three days. The frequency of GFP positive Raji cells was analyzed by flow cytometry on a FACS Canto II device. The percentages of GFP positive Raji cells was plotted against the administered volumes of the supernatant. The formula of a linear regression curve was calculated for the linear part of the graph using GraphPad Prism. The titer defined as “Green Raji Units/ml” was calculated as follows with **a** and **b** being given by the linear regression formula.

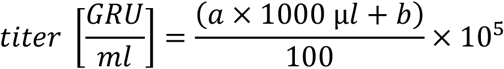

#### EBV mutagenesis

For EBV mutagenesis, we used 6008 EBV-BAC DNA which is based on B95.8 and contains all 44 miRNA (Pich *et al*., 2019). 6008 was provided by W. Hammerschmidt and used to introduce a C-terminal HA-tag in EBNA2 resulting in XZ143-EBV_HA-EBNA2 (EBVwt) (Zhang *et al*., 2021). This EBV genome (XZ143) was then used to delete the α1-helix in EBNA2 resulting in SB161-EBV_HA-EBNA2Δα1 (EBVΔα1). Recombinant EBV strains used in this study were generated by a two-step selection protocol using the λ-prophage-based heat inducible Red recombination system expressed in E.coli strain SW105 (Warming *et al*., 2005; Wang *et al*., 2009; Pich *et al*., 2019). Preparation of recombination-competent, electrocompetent SW105 bacteria and the electroporation of those was based on a protocol published on https://labnodes.vanderbilt.edu/resource/view/id/3165/collection_id/32/community_id/8.

In the first step, a rpsL/Kan expression cassette was amplified from p6012 template by PCR. 50 nt flanking regions homologous to the respective EBNA2 region were included. The resulting PCR product with the rpsL/Kan cassette was electroporated into SW105 carrying 6008 EBV-BAC and inserted into the specific EBNA2 region by homologous recombination. SW105 with successful recombination were selected with kanamycin (30μg/ml). As a second step, a synthetic DNA fragment carrying the desired mutation flanked by 300nt of the genomic viral sequence was used to replace the rpsL/Kan cassette by homologous recombination to generate the final mutant EBV plasmid. SW105 with successful recombination were selected with streptomycin (1mg/ml) and chloramphenicol (12.5 μg/ml).

#### EBV-BAC purification from bacterial cultures

Supercoiled EBV-BAC DNA was purified by CsCl_2_-EtBr density gradient centrifugation. 400 ml LB medium containing 15 μg/ml chloramphenicol were inoculated with a fresh single colony of SW105 bacteria carrying EBV-BAC of interest and cultivated at 32 °C shaking overnight. The bacteria were pelleted at 4000 rpm, 30 min, 4 °C. The BAC DNA was extracted using the NuceloBond™ Xtra BAC kit (Machery-Nagel) and according to the manufacturer’s instructions. The DNA was dissolved in 400 μl TE buffer overnight at 4 °C. The next day, 1.6 g CsCl_2_ was added to the solution, carefully dissolved and transferred to a 11.5 ml Ultracrimp tube (Sorvall).

The tube was filled up with 1.55 g/ml CsCl_2_ solution and 200 μl 1% EtBr. The sealed tube was ultracentrifuged at 35,000 rpm, 20 °C for 3 days. The lower band containing supercoiled DNA was extracted with a large gauge veterinary needle and transferred to 15 ml Falcon. EtBr was removed by isobutanol solvent extraction followed by a dialysis in 2 l TE at 4 °C overnight. The DNA concentration was determined by Qubit fluorometric quantitation.

#### Whole cell protein lysate

10^7^ cells were harvested, washed once with PBS and lysed with 200 μl NP-40 lysis buffer (1 % NP-40, 150 mM NaCl, 10 mM Tris-HCl pH7.4, EDTA pH8, 3 % Glycerol, 1x proteinase inhibitor cocktail (PIC, Roche), 1x PhosStop (Sigma-Aldrich)). The mix was incubated at 4 °C for one hour. Cell debris was pelleted at 16000 g, 4 °C for 15 min. Cleared lysate was transferred to a new tube. 25 μg of total protein were analyzed by SDS-PAGE and Western blot. Protein lysates were stored at −80 °C.

#### Purification of GST-tagged proteins

The purification of GST-tagged proteins was performed as previously described (Glaser *et al*., 2017). *E.coli* strain BL21 was transformed with the expression plasmids pET28_GST-END, pET28_GST-ENDΔα1, pET28_GST-END-H15A or pGEX-4T2 (GST only). Bacteria were cultured in 400 ml of LB medium with antibiotics at 37°C until an OD of 0.5 – 0.7 was reached. Due to a smaller yield, 800 ml LB culture were prepared for the expression of GST-ENDΔα1. The protein expression was induced with 1 mM IPTG for three hours at 30°C, shaking. After induction, bacteria were harvested at 4000 g, 20 min, 4 °C and resuspended in 20 ml ice cold binding buffer (25 mM HEPES, pH 7.6, 0.1 mM EDTA, pH 8, 12.5 mM MgCl_2_, 10% Glycerol, 0.1% NP-40, 100 mM KCl, 1 mM PMSF, 1 mM DTT). Bacteria were lysed by sonication (Brandson Digital Sonifier® 250, 10x 10 sec on/1 min off, 10 % amplitude). Lysates were cleared by centrifugation at 48,000 g, 4°C for 20 min. Glutathione Sepharose 4B beads (GE Healthcare) were washed and resuspended in binding buffer to prepare a 50% slurry. To coat the beads with GST or GST-tagged proteins, 100 μl of the 50% slurry were incubated with 20 ml of cleared lysates for one hour, 4°C, and washed three times with 20 ml binding buffer.

#### Affinity capture assay with recombinant GST and cell lysate

This assay was performed as previously described (Glaser *et al*., 2017). 10^7^ DG75 cells were transfected with 10 μg EBF1 expression plasmid or empty vector controls. 24 h after transfection, cells were harvested and lysed in 500 μl NP-40 lysis buffer (50 mM HEPES, pH 7.6, 5 mM EDTA, pH 8, 150 mM NaCl, 0.1% NP-40, 1 mM PMSF) followed by sonication. Cell lysates were centrifuged for 15 min at 16,000 g, 4°C, and the protein concentration was measured by Bradford assay. To pull down EBF1, the supernatants were incubated with the GST or GST fusion protein coated beads for three hours at 4°C. Subsequently, beads were washed 5x with binding buffer (25 mM HEPES, pH 7.6, 0.1 mM EDTA, pH 8, 12.5 mM MgCl_2_, 10% Glycerol, 0.1% NP-40, 100 mM KCl, 1 mM PMSF, 1 mM DTT) and the protein complexes were eluted in 2x Lämmli buffer (4% SDS, 20% Glycerol, 120 mM Tris/HCl, pH 6.8, 5%β-mercaptoethanol, bromophenol-blue). Samples were analyzed by SDS-PAGE and Western blot.

#### Co-Immunoprecipitation (Co-IP)

10 μg of plasmids or vector controls were transfected into 10^7^ DG75^doxHA-EBNA2^, DG75^doxHA-EBNA2ΔEND^, DG75^doxHA-EBNA2-H15A^ or DG75^doxHAEBNA2Δα1^ cells by electroporation at 250 V, 950 μF in 250 μl reduced serum medium (OptiMEM, Gibco) using the GenePulser II (Biorad). The cells were cultured with 1 μg/ml doxycycline overnight to induce the expression of EBNA2. The Co-IP was performed as previously described (Glaser *et al*., 2017). In brief, whole cell lysate was prepared with 520 μl NP-40 lysis buffer. A small aliquot was used for the input samples. For one IP, whole cell lysates were incubated with 250 μl α-EBNA2 antibody (R3, hybridoma supernatant, HMGU) or α-HA antibody (3F10) overnight at 4 °C. Next, 60 μl of equilibrated 50 % Protein G bead slurry were added and incubated for 3 hours at 4 °C. The beads were washed extensively with lysis buffer. The protein complexes were eluted with 60 μl 2x Lämmli buffer and heated to 95 °C for 5 min. 1:4 diluted input and undiluted Co-IP samples were analyzed by SDS-PAGE and Western blot.

#### Western Blot

Proteins resolved by SDS-PAGE were transferred to PVDF membranes for 1 hour at 400 mA. The membranes were blocked with blocking buffer (5 % non-fat dried milk powder in TBS) for 30 min at RT. Primary antibodies listed in Table 9 in blocking buffer were incubated with the membrane at 4 °C for one hour or overnight. After washing 3x with PBS/T for 10 min, the secondary HRP-coupled antibody in blocking buffer was incubated with the membrane at 4 °C for one hour followed by 4x washing with PBS/T for 15 min and a final wash with PBS. Bound antibodies were detected with the Enhanced Chemiluminescence (ECL) system (GE Healthcare Amersham) according to the manufacturer’s instructions. Emitted light was detected with the Fusion FX (Vilber) device.

#### Chromatin Immunoprecipitation (ChIP)

ChIP-qPCR was performed in LCLs as previously described (Glaser *et al*., 2017) with minor changes. In brief, 2×10^7^ cells were harvested, washed with PBS and resuspended in 20 ml RPMI medium. Cross-linking was achieved by the addition of formaldehyde (final 1 %) and 7 min incubation at room temperature. Glycine was added to a final concentration of 125 mM. The cells were washed in PBS, followed by washing with ice-cold lysis buffer (10 mM Tris-HCl, pH 7.5; 10 mM NaCl, 3 mM MgCl_2_, 0.5% NP-40, 1x proteinase inhibitor cocktail (PIC, Roche)). The cells were resuspended in sonication buffer (50 mM Tris-HCl, pH 8.0, 5 mM EDTA, pH 8.0, 0.5% SDS, 0.5% Triton X-100, 0.05% sodium deoxycholate, 1x PIC) and the chromatin was sheared by 4 rounds of 10 min sonication (Bioruptor, 30 sec on/off). 250 μl of cleared lysate containing chromatin was used for one IP. 25 μl of chromatin was saved for the input sample. For one α-EBNA2 ChIP, a 100 μl mix of α-EBNA2 (R3), α-EBNA2 (1E6) and α-HA (3F10) or a matched isotype control (α-GST (6G9) + α-Dog-CD3) was added to the chromatin. For one α-EBF1 ChIP, 5 μg of α-EBF1 (ABE1294) or a normal rabbit IgG (PP64B) isotype control was added to the chromatin. The chromatin-antibody solution was incubated overnight at 4 °C rotating. Protein A beads were used for the α-EBF1 ChIP and Protein G beads were used for the α-EBNA2 ChIP. The beads were pre-blocked with 500 μg/ml salmon testes DNA overnight at 4 °C. The beads were equilibrated with dilution buffer (12.5 mM Tri-HCl, pH 8.0, 187.5 mM NaCl, 1.25 mM EDTA, pH 8.0, 1.125% Triton X-100, 1 x PIC), 100 μl of a 50 % slurry was added to the IP and incubated at 4 °C for 3 hours. The beads were washed with: **2x Wash Buffer I** (20 mM Tris-HCl, pH 8.0, 2 mM EDTA, pH 8.0, 1% Triton X-100, 150 mM NaCl, 0.1% SDS, 1x PIC), **1x Wash Buffer II** (20 mM Tris-HCl, pH 8.0, 2 mM EDTA, pH 8.0, 1% Triton X-100, 500 mM NaCl, 0.1% SDS, 1x PIC), **1x Wash Buffer III** (10 mM Tris-HCl, pH 8.0, 1 mM EDTA, pH 8.0, 250 mM LiCl, 1% NP-40, 1% sodium deoxycholate, 1x PIC) for 5 min, rotating, and **2x with TE** (10 mM Tris-HCl, pH 8.0, 1 mM EDTA, pH 8.0) for 1 min. Chromatin was eluted with 2x 150 μl elution buffer (25 mM Tris-HCl, pH 7.5, 10 mM EDTA, pH 8.0, 1% SDS) and 15 min incubation at 65 °C. To reverse the cross-link, Proteinase K (final 1.5 mg/ml, Roche) was added to the input and ChIP samples, incubated for 1 hour at 42 °C and overnight at 65 °C. Reverse cross-linked chromatin was purified using the NucleoSpin™ Gel and PCR Clean-up kit (Macherey-Nagel) according to the manufacturer’s instructions. Purified ChIP and input samples were analyzed by qPCR.

#### cDNA preparation

RNA was isolated from 1×10^7^ cells with the RNeasy® Mini Kit (Qiagen) according to the manufacturer’s instructions. β-mercaptoethanol in a final concentration of 134 mM was added to the lysis buffer. Cells were lysed in 600 μl lysis buffer. QIAshredder columns (Qiagen) were used according to the manufacturer’s instructions. Genomic DNA was removed with the RNase-free DNase set (Qiagen) according to the manufacturer’s instructions. Purified RNA was eluted in 30-50 μl RNase-free water and the concentration was measured using the Qubit RNA HS kit according to the instructions. SuperScript™IV First-Strand cDNA Synthesis kit (Thermofisher) was used to transcribe 2-4 μg isolated RNA into cDNA according to the manufacturer’s instructions. Random hexamers were used as primers. A reaction without reverse transcriptase was used as a negative control. 1/40 of cDNA (50 ng of input RNA) was used for qPCR.

#### Quantitative real time PCR (qPCR)

The quantification of cDNA obtained from reverse transcribed RNA and DNA recovered from ChIP experiments was done with the Roche LightCycler480 II device and LightCycler 480 SYBR Green I Master (Roche) reagent according to the manufacturer’s instructions. The qPCR was carried out as previously described (Harth-Hertle *et al*., 2013). Amplification was performed at 63 °C. 10 μl per reaction were pipetted into a 96-well plate. 2 μl of the sample were added to each well. The master mix contained 5 μl LightCycler 480 SYBR Green I Master, 1 μl of 5 μM forward primer, 1 μl of 5 μM reverse primer, 1 μl PCR grade H_2_O. Standard curves with dilutions of defined amounts of the respective PCR product or chromatin were established and applied to account for varying primer efficiencies. Two replicates for each cDNA, ChIP target and standard dilution were cycled in parallel. The relative quantification of cDNA was normalized to RNA polymerase II signals applying the Roche LightCycler 480 E-Method, which is correcting for primer efficiencies.

The percentage of input for ChIP-qPCR signals was calculated for each target as (DNA from specific IP/ DNA input) x 100. Mann-Whitney test or Welch’s t-test were applied to test for differences.

#### RNA and cDNA library preparation for RNA-sequencing

RNA sequencing was performed using the prime-seq method based on the scRNA-seq method SCRB-seq (Bagnoli *et al*., 2018; Janjic *et al*., 2021). 10,000 cells per sample were sorted into a 96 well plate containing 50 μl RLT plus Lysis Buffer (Qiagen) supplemented with 1% 2-mercaptoethanol and frozen at −80 °C immediately. To assess how much RNA was extracted, replicate samples were pooled, prime-seq was performed on each pool individually and the cDNA amplification was performed as a qPCR using SYBR green. The input volume for the cDNA library preparation was adjusted accordingly.

**Table 10.** Volumes used for cDNA library preparation

| **Sample (6 replicates each)** | **Infection** | **Volume cell lysate [µl]** |
| --- | --- | --- |
| no_0 | non-infected | 50 |
| mt_1 | EBVΔα1 | 10 |
| mt_2 | EBVΔα1 | 50 |
| mt_3 | EBVΔα1 | 50 |
| mt_4 | EBVΔα1 | 30 |
| wt_1 | EBVwt | 30 |
| wt_2 | EBVwt | 10 |
| wt_3 | EBVwt | 10 |
| wt_4 | EBVwt | 10 |

Prime-seq was performed according to protocol (https://www.protocols.io/view/prime-seq-s9veh66). In brief, lysates were first treated with Proteinase K (Ambion) followed by a beads clean-up (2:1 beads to lysate ratio) using solid phase reversible immobilization (SPRI) beads (GE Biotech). A subsequent on-bead DNAseI digest was performed to remove genomic DNA, followed by another SPRI bead clean-up and reverse transcription. Reverse transcription (RT) was performed using Maxima H-RT enzyme (Thermo Fisher Scientific) with well specific barcoded oligo dT primers and a template switch oligo (TSO). After the RT, the barcoded first strand cDNA was pooled and cleaned up using magnetic beads. Remaining oligo dT primers were removed using exonucleaseI digest. Subsequently, cDNA was PCR amplified with Kapa Hifi DNA polymerase (Roche), quantified using picogreen dye (Thermo Fisher Scientific) and quality control using capillary gel electrophoresis (Agilent Bioanalyzer 2100). Sequencing libraries were generated with the NEB NEXT Ultra II FS kit by using a custom adapter oligo and a primer, specific for the cDNA 3’-end. Fragments between 300 bp and 500 bp were selected using double size selection with SPRIselect beads (Beckman Coulter). Final libraries were quantified and quality controlled again with the Bioanalyzer 2100. Paired End sequencing was performed as 28 bp read 1 and 50 bp read 2 with an 8 bp index read on an Illumina HiSeq 1500 instrument. In total 1.5 lanes of a High Out flow cell were sequenced, amounting to 465.9 Mio reads. Raw sequencing data was processed using the zUMIs pipeline (Parekh *et al*., 2018) with mapping to the reference genome using STAR 2.7 (Dobin *et al*., 2013). As a reference genome hg38 was used with Gencode annotation (v 35) concatenated with the EBV genomes Akata and p6008 (manually assembled, provided by W. Hammerschmidt) including annotation retrieved from (https://github.com/flemingtonlab/public/blob/master/annotation/chrEBV_Akata_inverted_refined_genes_annotation_cleaned.gtf)

#### Quality control and data normalization of RNA sequencing data

To initially assess the processed RNA sequencing data, the library size, number of detected cellular and viral genes and the fraction of mitochondrial genes was computed (Figure S4). For subsequent analyses, samples with less than 10,000 detected genes and a library size of less than 140,000 counts (one sample of each the day 0 non-infected, day 2 and day 3 EBVΔα1 infected condition: 14-no-0, 14-a1-2, 15-a1-3) were excluded. Data was normalized using size factors (Anders and Huber, 2010), which accounts for sequencing depth and RNA content.

#### Principal component analysis (PCA)

A principal component analysis (PCA) was performed on the log-transformed expression matrix of all protein-coding genes of all samples, with 1 added as a pseudo count.

#### Differential gene expression (DGE) analysis

Genes with a low expression (counts per million <10) were excluded. The DGE analysis was performed using the DESeq2 R package. Differentially expressed genes with an FDR < 0.1 and a fold change ≥ 2 were selected for the cluster analysis. Differentially expressed, protein coding genes with an FDR < 0.01 were selected for the gene set enrichment analysis (GSEA).

#### Gene set enrichment analysis (GSEA)

For the GSEA, the program was downloaded from https://www.gsea-msigdb.org as a desktop application. The hallmark gene sets were selected, and normalized expression data was used for the analysis. Enriched gene sets with an FDR <0.05 were considered significant.

#### Cluster analysis

To classify the patterns of gene expression along the infection time course, we applied a clustering algorithm to all DE genes in the EBVwt and EBVΔα1 infected cells separately. First, we calculated a distance matrix between genes as 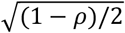, where *ρ* is the Spearman’s correlation coefficient between pairs of genes across all samples in a given condition. Hierarchical clustering was performed on this distance matrix (“hclust” function in R, with the “average” aggregation method), followed by the dynamic hybrid cut algorithm to estimate the number of clusters (dynamicTreeCut package v1.63-1, with minimum cluster size of 500 and “deepSplit” parameter equal to 1). We identified 5 clusters in EBVwt and 4 in EBVΔα1 (Figure S5 B-J) with the “clust.stats” function from the “fpc” R package (version 2.1-11.1).

#### Gene Ontology (GO) Analysis

The GO term analysis was performed with the online tool GOrilla (Eden *et al*., 2007, 2009) in order to identify biological processes enriched in each cluster. The analysis was performed with a list of genes identified for each cluster and a background list of genes that included all genes that were tested for differential expression. The p-value threshold for the GO term analysis was set to 10^-3. The online tool REVIGO was used to reduce redundancy of identified GO terms and the analyses were performed with settings defining medium similarity, the homo sapiens databank and SimRel to calculate semantic similarity (Supek *et al*., 2011).

#### Data and code availability

The raw and processed sequencing data will be available from Array express. All data were analyzed with standard programs and packages, as detailed above. Code is available on request.

## REFERENCES

Akkaya, M. and Pierce, S. K. (2019) ‘From zero to sixty and back to zero again: the metabolic life of B cells’, Current Opinion in Immunology, 57, pp. 1–7. doi: 10.1016/j.coi.2018.09.019.

Anders, S. and Huber, W. (2010) ‘Differential expression analysis for sequence count data’, Genome Biology. BioMed Central, 11(10), p. R106. doi: 10.1186/gb-2010-11-10-r106.

Arnett, K. L. et al. (2010) ‘Structural and mechanistic insights into cooperative assembly of dimeric Notch transcription complexes’, Nature Structural and Molecular Biology. Nature Publishing Group, 17(11), pp. 1312–1317. doi: 10.1038/nsmb.1938.

Bagnoli, J. W. et al. (2018) ‘Sensitive and powerful single-cell RNA sequencing using mcSCRB-seq’, Nature Communications. Nature Publishing Group, 9(1), pp. 1–8. doi: 10.1038/s41467-018-05347-6.

Banchereau, J. et al. (1990) ‘Long-term human B cell lines dependent on interleukin-4 and antibody to CD40’, Science. Science, 251(4989), pp. 70–72. doi: 10.1126/science.1702555.

Ben-bassats, H. et al. (1977) ‘Establishment in continuous culture of a new type of lymphocyte from a “burkitt-like” malignant lymphoma (line d.g.-75)’, International Journal of Cancer, 19(1), pp. 27–33. doi: 10.1002/ijc.2910190105.

Bohle, V. et al. (2013) ‘Role of early B-cell factor 1 (EBF1) in Hodgkin lymphoma’, Leukemia. Leukemia, 27(3), pp. 671–679. doi: 10.1038/leu.2012.280.

Boller, S. et al. (2016) ‘Pioneering Activity of the C-Terminal Domain of EBF1 Shapes the Chromatin Landscape for B Cell Programming’, Immunity, 44(3), pp. 527–541. doi: 10.1016/j.immuni.2016.02.021.

Boller, S., Li, R. and Grosschedl, R. (2018) ‘Defining B Cell Chromatin: Lessons from EBF1’, Trends in Genetics. Elsevier Ltd, xx, pp. 1–13. doi: 10.1016/j.tig.2017.12.014.

Bullerwell, C. E. et al. (2021) ‘EBF1 drives hallmark B cell gene expression by enabling the interaction of PAX5 with the MLL H3K4 methyltransferase complex’, Scientific Reports. Springer Science and Business Media LLC, 11(1). doi: 10.1038/s41598-021-81000-5.

Cohen, J. I. et al. (1989) ‘Epstein-Barr virus nuclear protein 2 is a key determinant of lymphocyte transformation’, Proceedings of the National Academy of Sciences of the United States of America. Proc Natl Acad Sci U S A, 86(23), pp. 9558–9562. doi: 10.1073/pnas.86.23.9558.

Cohen, J. I., Wang, F. and Kieff, E. (1991) ‘Epstein-Barr virus nuclear protein 2 mutations define essential domains for transformation and transactivation’, J Virol, 65(5), pp. 2545–2554.

Dirmeier, U. et al. (2005) ‘Latent membrane protein 1 of Epstein-Barr virus coordinately regulates proliferation with control of apoptosis’, Oncogene. Oncogene, 24(10), pp. 1711–1717. doi: 10.1038/sj.onc.1208367.

Dobin, A. et al. (2013) ‘STAR: Ultrafast universal RNA-seq aligner’, Bioinformatics. Bioinformatics, 29(1), pp. 15–21. doi: 10.1093/bioinformatics/bts635.

Dong, Y. et al. (2020) ‘Regulation of cancer cell metabolism: oncogenic MYC in the driver’s seat’, Signal Transduction and Targeted Therapy. doi: 10.1038/s41392-020-00235-2.

Eden, E. et al. (2007) ‘Discovering Motifs in Ranked Lists of DNA Sequences’, PLOS Computational Biology. Public Library of Science, 3(3), p. e39. doi: 10.1371/JOURNAL.PCBI.0030039.

Eden, E. et al. (2009) ‘GOrilla: a tool for discovery and visualization of enriched GO terms in ranked gene lists’, BMC Bioinformatics 2009 10:1. BioMed Central, 10(1), pp. 1–7. doi: 10.1186/1471-2105-10-48.

Egawa, T. and Bhattacharya, D. (2019) ‘Regulation of metabolic supply and demand during B cell activation and subsequent differentiation’, Current opinion in immunology. NIH Public Access, 57, p. 8. doi: 10.1016/J.COI.2018.10.003.

Elgueta, R. et al. (2009) ‘Molecular mechanism and function of CD40/CD40L engagement in the immune system’, Immunological Reviews. NIH Public Access, pp. 152–172. doi: 10.1111/j.1600-065X.2009.00782.x.

Friberg, A. et al. (2015) ‘The EBNA-2 N-Terminal Transactivation Domain Folds into a Dimeric Structure Required for Target Gene Activation’, PLoS Pathogens, 11(5), pp. 1–24. doi: 10.1371/journal.ppat.1004910.

Gao, H. et al. (2009) ‘Opposing effects of SWI/SNF and Mi-2/NuRD chromatin remodeling complexes on epigenetic reprogramming by EBF and Pax5’, Proceedings of the National Academy of Sciences, 106(27), pp. 11258–11263.

Glaser, L. V. et al. (2017) ‘EBF1 binds to EBNA2 and promotes the assembly of EBNA2 chromatin complexes in B cells’, PLoS Pathogens, 13(10), pp. 1–30. doi: 10.1371/journal.ppat.1006664.

Graham, F. L. et al. (1977) Characteristics of a Human Cell Line Transformed by DNA from Human Adenovirus Type 5, J. gen. Virol.

Grossman, S. R. et al. (1994) ‘The Epstein-Barr virus nuclear antigen 2 transactivator is directed to response elements by the Jκ recombination signal binding protein’, Proceedings of the National Academy of Sciences of the United States of America. Proc Natl Acad Sci U S A, 91(16), pp. 7568–7572. doi: 10.1073/pnas.91.16.7568.

Györy, I. et al. (2012) ‘Transcription factor EBf1 regulates differentiation stage-specific signaling, proliferation, and survival of B cells’, Genes and Development, 26(7), pp. 668–682. doi: 10.1101/gad.187328.112.

Hagman, J., Travis, A. and Grosschedl, R. (1991) ‘A novel lineage-specific nuclear factor regulates mb-1 gene transcription at the early stages of B cell differentiation’, EMBO Journal, 10(11), pp. 3409–3417.

Hammerschmidt, W. and Sugden, B. (1989) ‘Genetic analysis of immortalizing functions of Epstein-Barr virus in human B lymphocytes’, Nature, 340(6232), pp. 393–397.

Harth-Hertle, M. L. et al. (2013) ‘Inactivation of Intergenic Enhancers by EBNA3A Initiates and Maintains Polycomb Signatures across a Chromatin Domain Encoding CXCL10 and CXCL9’, PLoS Pathogens. Edited by P. M. Lieberman, 9(9), p. e1003638. doi: 10.1371/journal.ppat.1003638.

Icard, P. et al. (2019) ‘Interconnection between Metabolism and Cell Cycle in Cancer’, Trends in Biochemical Sciences. Elsevier Current Trends, 44(6), pp. 490–501. doi: 10.1016/J.TIBS.2018.12.007.

Janjic, A. et al. (2021) ‘Prime-seq, efficient and powerful bulk RNA-sequencing’, bioRxiv. Cold Spring Harbor Laboratory, p. 2021.09.27.459575. doi: 10.1101/2021.09.27.459575.

Jiang, S. et al. (2017) ‘The Epstein-Barr Virus Regulome in Lymphoblastoid Cells’, Cell Host & Microbe. Cell Press, 22(4), pp. 561–573.e4. doi: 10.1016/J.CHOM.2017.09.001.

Jiang, S. et al. (2018) ‘CRISPR/Cas9-Mediated Genome Editing in Epstein-Barr Virus-Transformed Lymphoblastoid B-Cell Lines’, Current Protocols in Molecular Biology, (January), pp. 31.12.1–31.12.23. doi: 10.1002/cpmb.51.

Johannsen, E. et al. (1995) ‘Epstein-Barr virus nuclear protein 2 transactivation of the latent membrane protein 1 promoter is mediated by J kappa and PU.1’, Journal of Virology. American Society for Microbiology, 69(1), pp. 253–262. doi: 10.1128/jvi.69.1.253-262.1995.

Kaiser, C. et al. (1999) ‘The Proto-Oncogene c-myc Is a Direct Target Gene of Epstein-Barr Virus Nuclear Antigen 2’, Journal of Virology. American Society for Microbiology, 73(5), pp. 4481–4484. doi: 10.1128/jvi.73.5.4481-4484.1999.

Kempkes, B., Spitkovsky, D., et al. (1995) ‘B-cell proliferation and induction of early G1-regulating proteins by Epstein-Barr virus mutants conditional for EBNA2.’, The EMBO Journal. European Molecular Biology Organization, 14(1), p. 88.

Kempkes, B., Pich, D., et al. (1995) ‘Immortalization of human primary B lymphocytes in vitro with DNA’, Proc Natl Acad Sci U S A, 92(13), pp. 5875–5879.

Kempkes, B. and Ling, P. D. (2015) ‘EBNA2 and its coactivator EBNA-LP’, in Current Topics in Microbiology and Immunology. Springer Verlag, pp. 35–59. doi: 10.1007/978-3-319-22834-1_2.

Laux, G. et al. (1994) ‘The Spi-1/PU.1 and Spi-B ets family transcription factors and the recombination signal binding protein RBP-J kappa interact with an Epstein-Barr virus nuclear antigen 2 responsive cis-element.’, The EMBO Journal. Wiley-VCH Verlag, 13(23), pp. 5624–5632. doi: 10.1002/j.1460-2075.1994.tb06900.x.

Liberzon, A. et al. (2015) ‘The Molecular Signatures Database Hallmark Gene Set Collection’, Cell Systems. Cell Press, 1(6), pp. 417–425. doi: 10.1016/j.cels.2015.12.004.

Ling, P. D., Rawlins, D. R. and Hayward, S. D. (1993) ‘The Epstein-Barr virus immortalizing protein EBNA-2 is targeted to DNA by a cellular enhancer-binding protein.’, Proceedings of the National Academy of Sciences of the United States of America, 90(20), pp. 9237–41. doi: 10.1073/pnas.90.20.9237.

Lu, F. et al. (2016) ‘EBNA2 Drives Formation of New Chromosome Binding Sites and Target Genes for B-Cell Master Regulatory Transcription Factors RBP-jκ and EBF1’, PLOS Pathogens. Edited by E. K. Flemington, 12(1), p. e1005339. doi: 10.1371/journal.ppat.1005339.

McClellan, M. J. et al. (2013) ‘Modulation of Enhancer Looping and Differential Gene Targeting by Epstein-Barr Virus Transcription Factors Directs Cellular Reprogramming’, PLOS Pathogens. Public Library of Science, 9(9), p. e1003636. doi: 10.1371/JOURNAL.PPAT.1003636.

Mrozek-Gorska, P. et al. (2019) ‘Epstein-Barr virus reprograms human B lymphocytes immediately in the prelatent phase of infection.’, Proceedings of the National Academy of Sciences of the United States of America. National Academy of Sciences. doi: 10.1073/pnas.1901314116.

Münz, C. (2019) ‘Latency and lytic replication in Epstein–Barr virus-associated oncogenesis’, Nature Reviews Microbiology. Nature Publishing Group, pp. 691–700. doi: 10.1038/s41579-019-0249-7.

Murata, T. et al. (2016) ‘Induction of Epstein-Barr Virus Oncoprotein LMP1 by Transcription Factors AP-2 and Early B Cell Factor’, J Virol, 90(8), pp. 3873–3889. doi: 10.1128/JVI.03227-15.

Nie, Z. et al. (2012) ‘c-Myc Is a Universal Amplifier of Expressed Genes in Lymphocytes and Embryonic Stem Cells’, Cell. Elsevier, 151(1), pp. 68–79. doi: 10.1016/j.cell.2012.08.033.

Parekh, S. et al. (2018) ‘zUMIs - A fast and flexible pipeline to process RNA sequencing data with UMIs’, GigaScience. Gigascience. doi: 10.1093/gigascience/giy059.

Pich, D. et al. (2019) ‘First Days in the Life of Naive Human B Lymphocytes Infected with Epstein-Barr Virus’, mBio. Edited by R. F. Ambinder and D. E. Griffin, 10(5), p. 17. doi: 10.1128/mBio.01723-19.

Polack, A. et al. (1996) ‘c-myc activation renders proliferation of Epstein-Barr virus (EBV)-transformed cells independent of EBV nuclear antigen 2 and latent membrane protein 1’, Proc Natl Acad Sci U S A, 93(19), pp. 10411–10416.

Pulvertaft, R. J. V. (1964) ‘CYTOLOGY OF BURKITT’S TUMOUR (AFRICAN LYMPHOMA)’, The Lancet, 283(7327), pp. 238–240. doi: 10.1016/S0140-6736(64)92345-1.

Shannon-Lowe, C., Rickinson, A. B. and Bell, A. I. (2017) ‘Epstein-barr virus-associated lymphomas’, Philosophical Transactions of the Royal Society B: Biological Sciences. Philos Trans R Soc Lond B Biol Sci. doi: 10.1098/rstb.2016.0271.

Sigvardsson, M. (2018) ‘Molecular Regulation of Differentiation in Early B-Lymphocyte Development’, International Journal of Molecular Sciences, 19(7), p. 1928. doi: 10.3390/ijms19071928.

Somasundaram, R. et al. (2021) ‘EBF1 and PAX5 control pro-B cell expansion via opposing regulation of the Myc gene’, Blood. American Society of Hematology, 137(22), pp. 3037–3049. doi: 10.1182/BLOOD.2020009564.

Strobl, L. J. et al. (1997) ‘Both Epstein-Barr viral nuclear antigen 2 (EBNA2) and activated notch 1 transactivate genes by interacting with the cellular protein RBP-Jκ’, Immunobiology. Urban & Fischer, 198(1–3), pp. 299–306. doi: 10.1016/S0171-2985(97)80050-2.

Supek, F. et al. (2011) ‘REVIGO Summarizes and Visualizes Long Lists of Gene Ontology Terms’, PLOS ONE. Public Library of Science, 6(7), p. e21800. doi: 10.1371/JOURNAL.PONE.0021800.

Thorley-Lawson, D. A. (2001) ‘Epstein-Barr virus: Exploiting the immune system’, Nature Reviews Immunology. European Association for Cardio-Thoracic Surgery, pp. 75–82. doi: 10.1038/35095584.

Vilagos, B. et al. (2012) ‘Essential role of EBF1 in the generation and function of distinct mature B cell types’, Journal of Experimental Medicine, 209(4), pp. 775–792. doi: 10.1084/jem.20112422.

Waltzer, L. et al. (1994) ‘The human J kappa recombination signal sequence binding protein (RBP-J kappa) targets the Epstein-Barr virus EBNA2 protein to its DNA responsive elements’, Embo J, 13(23), pp. 5633–5638.

Wang, S. et al. (2009) ‘A new positive/negative selection scheme for precise BAC recombineering’, Molecular Biotechnology, 42, pp. 110–116. doi: 10.1007/s12033-009-9142-3.

Wang, Y. et al. (2020) ‘A Prion-like Domain in Transcription Factor EBF1 Promotes Phase Separation and Enables B Cell Programming of Progenitor Chromatin’, Immunity. Cell Press, 53(6), pp. 1151–1167.e6. doi: 10.1016/j.immuni.2020.10.009.

Warming, S. et al. (2005) ‘Simple and highly efficient BAC recombineering using galK selection’, Nucleic Acids Res, 33(4). doi: 10.1093/nar/gni035.

Wiesner, Martina et al. (2008) ‘Conditional Immortalization of Human B Cells by CD40 Ligation’, PLoS ONE. Edited by J. Alberola-Ila. Public Library of Science, 3(1), p. e1464. doi: 10.1371/journal.pone.0001464.

Wiesner, M et al. (2008) ‘Conditional Immortalization of Human B Cells by CD40 Ligation’, PLoS ONE, 3(1), p. 1464. doi: 10.1371/journal.pone.0001464.

Wood, C. D. et al. (2016) ‘MYC activation and BCL2L11 silencing by a tumour virus through the large-scale reconfiguration of enhancer-promoter hubs’, eLife. eLife Sciences Publications, Ltd, 5(AUGUST). doi: 10.7554/ELIFE.18270.

Wu, D. Y. et al. (1996) ‘Epstein-Barr virus nuclear protein 2 (EBNA2) binds to a component of the human SNF-SWI complex, hSNF5/Ini1’, J Virol, 70(9), pp. 6020–8.

Yalamanchili, R. et al. (1994) ‘Genetic and biochemical evidence that EBNA 2 interaction with a 63-kDa cellular GTG-binding protein is essential for B lymphocyte growth transformation by EBV.’, Virology, pp. 634–41. doi: 10.1006/viro.1994.1578.

Zhang, X. et al. (2021) ‘PLK1-dependent phosphorylation restrains EBNA2 activity and lymphomagenesis in EBV-infected mice’, EMBO reports. EMBO Rep, 22(12). doi: 10.15252/EMBR.202153007.

Zhao, B. et al. (2011) ‘Epstein-Barr virus exploits intrinsic B-lymphocyte transcription programs to achieve immortal cell growth.’, Proceedings of the National Academy of Sciences of the United States of America, 108(36), pp. 14902–7. doi: 10.1073/pnas.1108892108.

Zhou, H. et al. (2015) ‘Epstein-barr virus oncoprotein super-enhancers control B cell growth’, Cell Host and Microbe. Cell Press, 17(2), pp. 205–216. doi: 10.1016/j.chom.2014.12.013

Zimber-Strobl, U. et al. (1994) ‘Epstein-Barr virus nuclear antigen 2 exerts its transactivating function through interaction with recombination signal binding protein RBP-J kappa, the homologue of Drosophila Suppressor of Hairless’, Embo J, 13(20), pp. 4973–4982.

Zimber-Strobl, U. and Strobl, L. J. (2001) ‘EBNA2 and Notch signalling in Epstein-Barr virus mediated immortalization of B lymphocytes’, Semin Cancer Biol, 11(6), pp. 423–34.

