## Supplemental figures 1-5 and tables 1-5 for "The EBNA2-EBF1 complex promotes oncogenic MYC expression levels and metabolic processes required for cell cycle progression of Epstein-Barr virus-infected B cells"

### SUPPLEMENTS

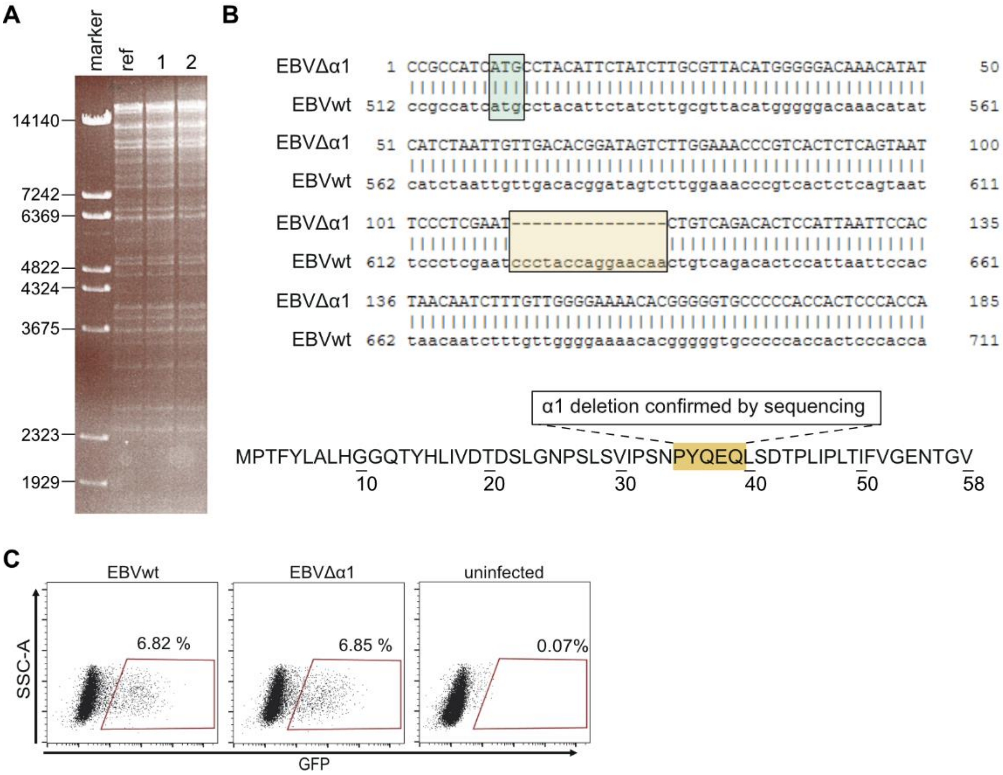


Figure S 1 Generating EBVΔα1 by recombineering. (A) Agarose gel of CsCl purified, KpnI digested EBV-BAC from EBVwt (XZ143, ref) and EBVΔα1 (SB161, 1 and 2), BstEII digested λ phage DNA as DNA marker (B) Alignment of the sequenced EBNA2 region in EBVΔα1 and EBVwt. Yellow box indicates α1-helix deletion. Green box indicates EBNA2 start codon. Amino acid sequence of the END domain of EBNA2 with yellow highlighting the α1 amino acid sequence. (C) Infection of Raji cells with 100 µl viral supernatants shows comparable infectivity of EBVwt and EBVΔα1.

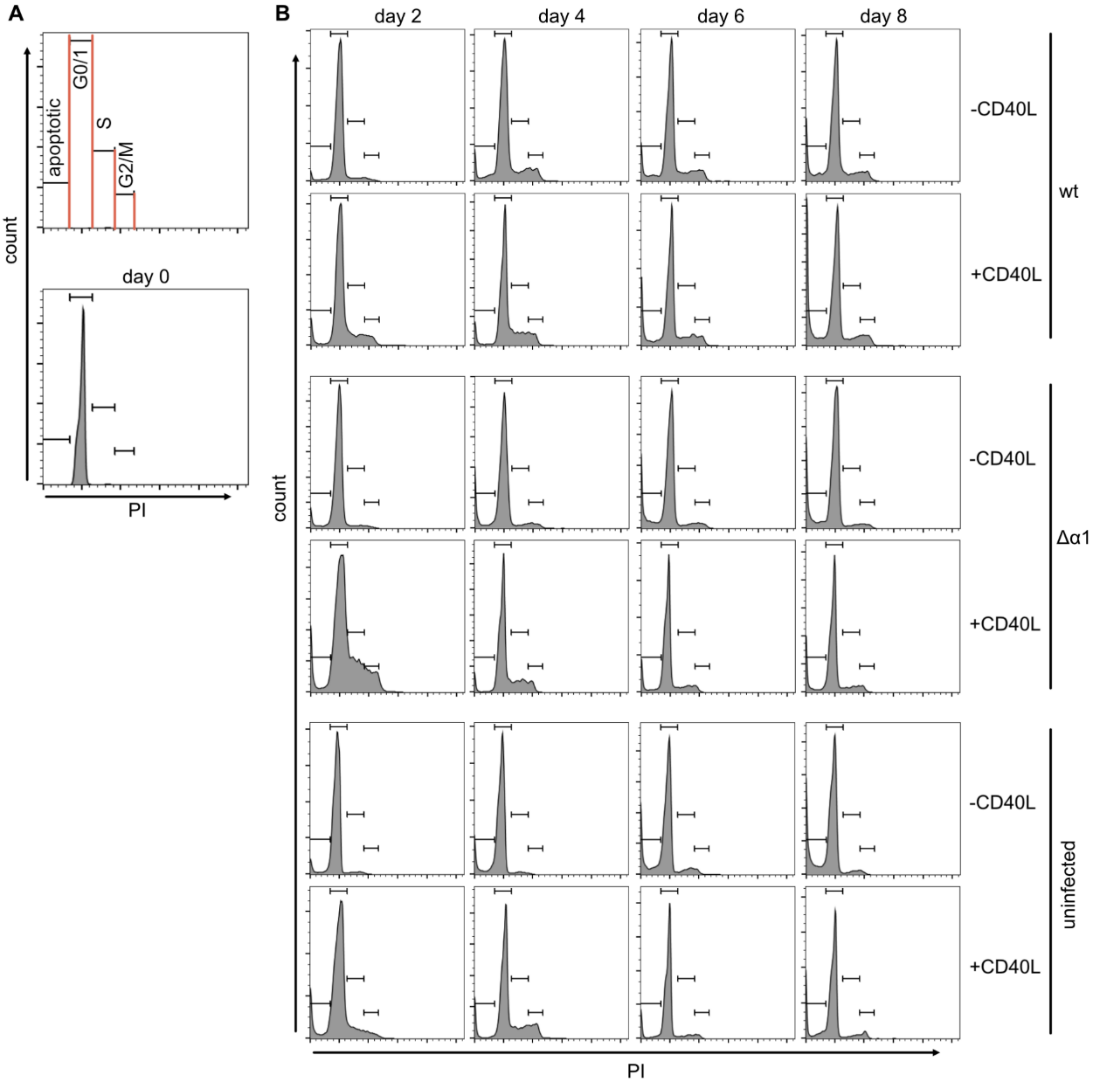


Figure S 2 Histograms of cell cycle analysis with propidium iodide (PI) (A) schematic gating of the cell cycle populations and one representative FACS plot for the day 0 non-infected sample, gated on lymphocytes and single cells. (B) PI histograms of one representative experiment showing B cells infected with EBVwt (wt), EBVΔα1 (Δα1) or non-infected B cells on day 2, 4, 6 and 8 post-infection. B cells were cultured without (-) CD40L expressing feeder cells or with (+) CD40L expressing feeder cells.

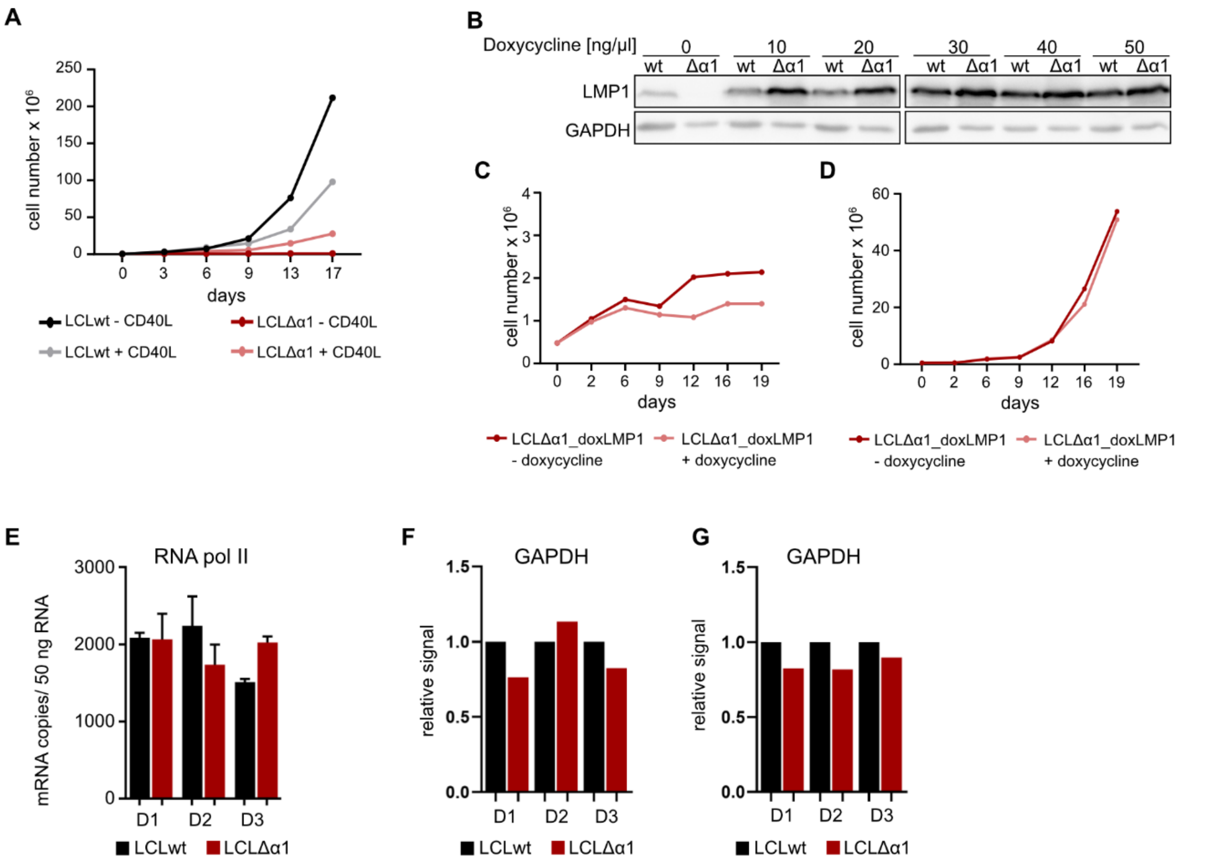


Figure S 3 Establishment of LCLΔα1 cultures on CD40L expressing feeder cells. (A) Growth curves of LCLwt and LCLΔα1 without (-) or with (+) CD40L feeder cells. $5 \times 10^5$ cells were seeded on day 0. The cell number was counted on the indicated time points. (B) LMP1 expression in LCLwt and LCLΔα1 transfected with the vector plasmid pRTS-HA-LMP1, which expresses LMP1 conditionally upon addition of doxycycline. GAPDH was used as a loading control. (C, D) Growth curves of LCLΔα1_doxLMP1 without (- doxycycline) and with ectopically expressed LMP1 (+ doxycycline) grown (C) without CD40L feeder cells or (D) with CD40L feeder cells. LMP1 expression was induced with 10 ng/ml doxycycline. $5 \times 10^5$ cells were seeded on day 0. Cell numbers were counted at the indicated time points. The mean of two replicates is plotted. (E) RT-qPCR quantification of RNA pol II transcripts in LCLwt and LCLΔα1 from three donors (D1, D2, D3). Error bars indicate standard deviation. (F, G) Quantified GAPDH signals from Western blots in figure 3D with (F) corresponding to the upper panel and (G) corresponding to the lower panel in figure 3D. The signal intensity was calculated with the Bio1D software and normalized to the GAPDH signals in LCLwt per donor and set to 1.

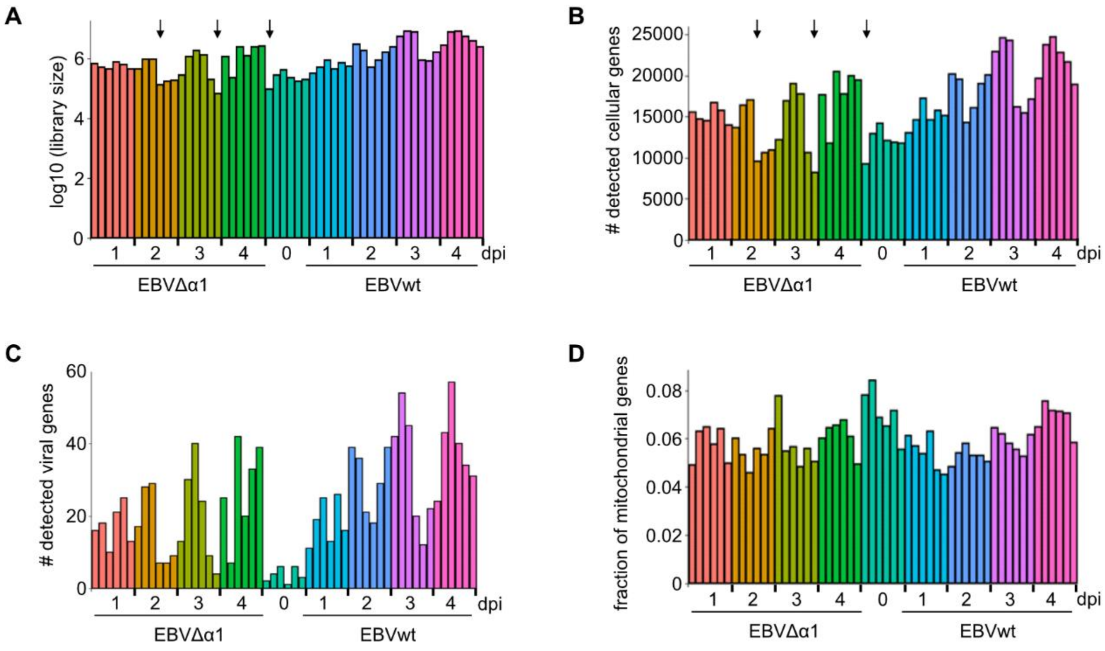


Figure S 4 Quality control of RNA sequencing samples. 6 biological replicates for each time point and infection condition were collected. (A) Log10 transformed size of sequenced libraries for all collected samples in infection conditions and at different days post-infection. (B) Number of detected cellular genes. (C) Number of detected viral genes. In total, 4920 reads were mapped to the viral genome. 3095 of these were mapped to EBNA2. (D) Fraction of mitochondrial genes detected among cellular genes. dpi – days post-infection, Arrows indicated samples that were excluded from the analysis.

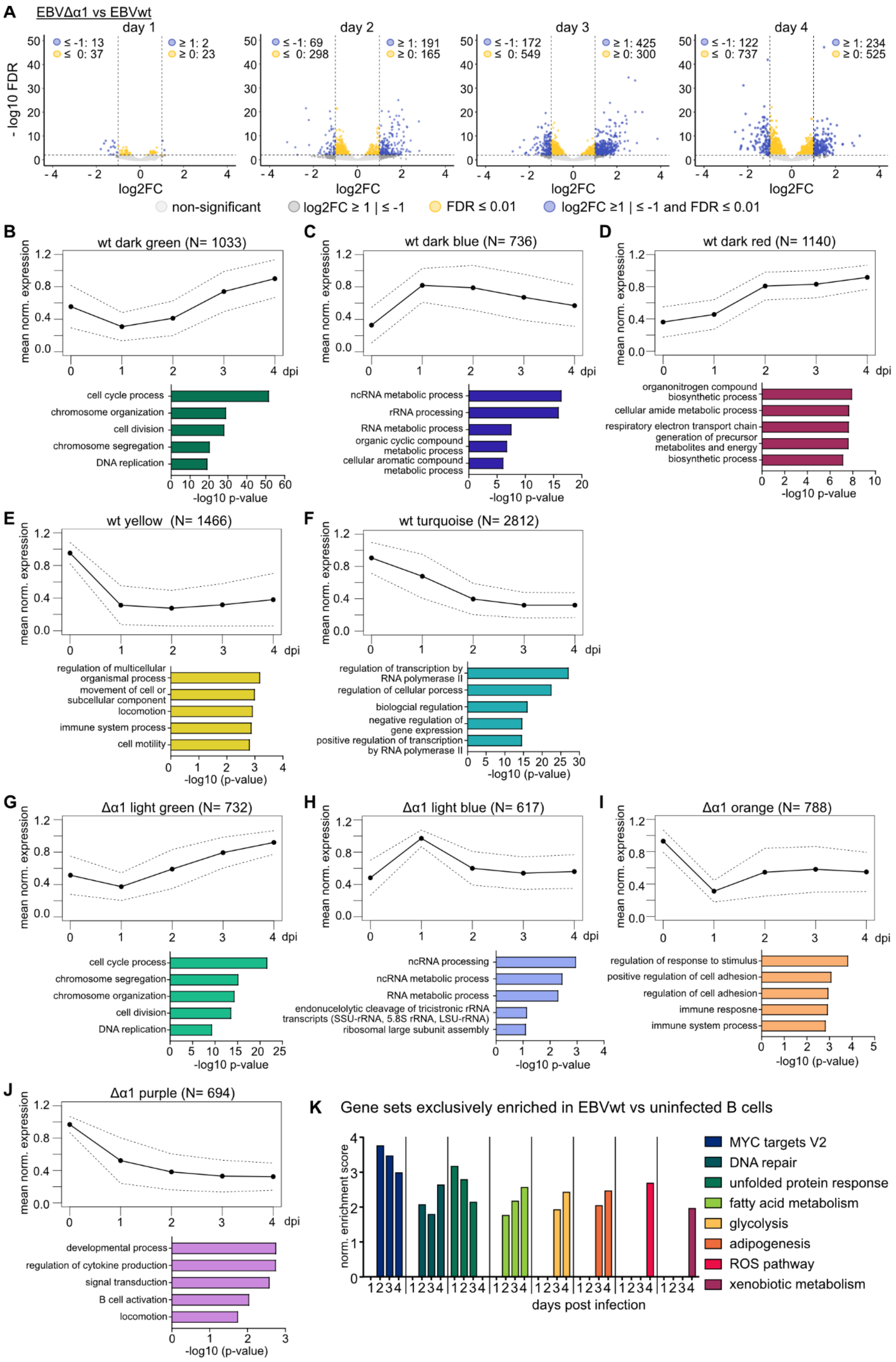
A
EBVΔα1 vs EBVwt
day 1
≤ -1: 13
≤ 0: 37
≥ 1: 2
≥ 0: 23
day 2
≤ -1: 69
≤ 0: 298
≥ 1: 191
≥ 0: 165
day 3
≤ -1: 172
≤ 0: 549
≥ 1: 425
≥ 0: 300
day 4
≤ -1: 122
≤ 0: 737
≥ 1: 234
≥ 0: 525
- log10 FDR
log2FC
non-significant
log2FC ≥ 1 | ≤ -1
FDR ≤ 0.01
log2FC ≥1 | ≤ -1 and FDR ≤ 0.01
B
wt dark green (N= 1033)
mean norm. expression
dpi
cell cycle process
chromosome organization
cell division
chromosome segregation
DNA replication
-log10 p-value
C
wt dark blue (N= 736)
ncRNA metabolic process
rRNA processing
RNA metabolic process
organic cyclic compound metabolic process
cellular aromatic compound metabolic process
-log10 p-value
D
wt dark red (N= 1140)
organonitrogen compound biosynthetic process
cellular amide metabolic process
respiratory electron transport chain
generation of precursor metabolites and energy
biosynthetic process
-log10 p-value
E
wt yellow (N= 1466)
regulation of multicellular organismal process
movement of cell or subcellular component
locomotion
immune system process
cell motility
-log10 (p-value)
F
wt turquoise (N= 2812)
regulation of transcription by RNA polymerase II
regulation of cellular porcess
biologcial regulation
negative regulation of gene expression
positive regulation of transcription by RNA polymerase II
-log10 (p-value)
G
Δα1 light green (N= 732)
cell cycle process
chromosome segregation
chromosome organization
cell division
DNA replication
-log10 p-value
H
Δα1 light blue (N= 617)
ncRNA processing
ncRNA metabolic process
RNA metabolic process
endonucelolytic cleavage of tricistronic rRNA transcripts (SSU-rRNA, 5.8S rRNA, LSU-rRNA)
ribosomal large subunit assembly
-log10 p-value
I
Δα1 orange (N= 788)
regulation of response to stimulus
positive regulation of cell adhesion
regulation of cell adhesion
immune resposne
immune system process
-log10 (p-value)
J
Δα1 purple (N= 694)
developmental process
regulation of cytokine production
signal transduction
B cell activation
locomotion
-log10 (p-value)
K
Gene sets exclusively enriched in EBVwt vs uninfected B cells
norm. enrichment score
days post infection
MYC targets V2
DNA repair
unfolded protein response
fatty acid metabolism
glycolysis
adipogenesis
ROS pathway
xenobiotic metabolism

Figure S 5 Time-resolved dynamic changes of gene expression patterns. (A) Direct comparison of EBVΔα1 and EBVwt infected B cells shown as differentially expressed (DE), protein coding genes in Volcano plots. Dotted lines indicate log2FC = 1 and FDR = 0.01. Cluster analyses of DE genes in (B-F) EBVwt and (G-J) EBVΔα1 infected B cells. The normalized mean expression is plotted with the standard deviation as dotted lines. N indicates the number of DE genes within each cluster. All genes from each cluster were used to perform the GO term analysis summarized in the bar graphs below. The top 5 GO terms according to the -log10 p-value are plotted. (K) Significantly enriched gene sets that are exclusively detected in EBVwt infected B cells. Differentially expressed, protein coding genes defined by the analysis EBVwt vs non-infected were included for this GSEA. Bar graphs showing the normalized enrichment score (NES) for each gene set at each day post-infection.

Table S 1 Corresponding to figure 2: Cell cycle distribution of EBVwt infected B cells analyzed by BrdU incorporation and 7-AAD staining. The mean was used to generate the plot in figure 2D. dpi – days post-infection

| **EBVwt** | | | | | |
|---|---|---|---|---|---|
| Donor 6 | | | | | |
| dpi | apoptotic | G0/1 | early S | S | G2/M |
| 0 | 0.10 | 90.74 | 1.87 | 5.02 | 2.26 |
| 2 | 0.30 | 70.50 | 19.06 | 7.91 | 2.23 |
| 4 | 2.96 | 52.10 | 17.78 | 18.92 | 8.25 |
| 6 | 2.82 | 66.96 | 9.69 | 13.82 | 6.70 |
| 8 | 5.19 | 75.16 | 6.99 | 7.29 | 5.38 |
| Donor 10 | | | | | |
| dpi | apoptotic | G0/1 | early S | S | G2/M |
| 0 | 0.15 | 91.10 | 2.52 | 2.61 | 3.62 |
| 2 | 2.56 | 73.49 | 14.74 | 6.75 | 2.47 |
| 4 | 4.89 | 67.91 | 4.43 | 11.44 | 11.33 |
| 6 | 4.37 | 78.04 | 3.06 | 6.81 | 7.72 |
| 8 | 4.85 | 82.82 | 2.54 | 3.49 | 6.30 |
| Donor 21 | | | | | |
| dpi | apoptotic | G0/1 | early S | S | G2/M |
| 0 | 1.09 | 90.58 | 1.89 | 3.54 | 2.89 |
| 2 | 3.32 | 66.66 | 23.09 | 5.28 | 1.64 |
| 4 | 5.63 | 76.39 | 6.66 | 6.35 | 4.98 |
| 6 | 8.62 | 75.31 | 7.75 | 4.08 | 4.23 |
| 8 | 11.86 | 75.37 | 4.59 | 1.93 | 6.25 |
| mean | | | | | |
| dpi | apoptotic | G0/1 | early S | S | G2/M |
| 0 | 0.45 | 90.81 | 2.09 | 3.73 | 2.92 |
| 2 | 2.06 | 70.20 | 18.98 | 6.64 | 2.11 |
| 4 | 4.50 | 65.56 | 9.58 | 12.18 | 8.18 |
| 6 | 5.28 | 73.44 | 6.84 | 8.23 | 6.21 |
| 8 | 7.31 | 77.82 | 4.69 | 4.21 | 5.98 |

Table S 2 Corresponding to figure 2: Cell cycle distribution of EBVΔα1 infected B cells analyzed by BrdU incorporation and 7-AAD staining. The mean was used to generate the plot in figure 2D. dpi – days post-infection

| **EBVΔα1** | | | | | |
|---|---|---|---|---|---|
| Donor 6 | | | | | |
| dpi | apoptotic | G0/1 | early S | S | G2/M |
| 0 | 0.10 | 90.74 | 1.87 | 5.02 | 2.26 |
| 2 | 0.25 | 75.00 | 20.26 | 3.32 | 1.18 |
| 4 | 4.38 | 62.04 | 25.00 | 6.39 | 2.18 |
| 6 | 7.46 | 62.11 | 19.72 | 6.18 | 4.53 |
| 8 | 17.40 | 54.47 | 20.64 | 5.43 | 2.06 |
| Donor 10 | | | | | |
| dpi | apoptotic | G0/1 | early S | S | G2/M |
| 0 | 0.15 | 91.10 | 2.52 | 2.61 | 3.62 |
| 2 | 2.93 | 76.15 | 15.44 | 3.64 | 1.84 |
| 4 | 6.61 | 77.40 | 7.55 | 2.90 | 5.54 |
| 6 | 9.11 | 70.80 | 11.85 | 3.35 | 4.89 |
| 8 | 10.84 | 74.53 | 8.86 | 1.57 | 4.21 |
| Donor 21 | | | | | |
| dpi | apoptotic | G0/1 | early S | S | G2/M |
| 0 | 1.09 | 90.58 | 1.89 | 3.54 | 2.89 |
| 2 | 3.62 | 68.66 | 20.83 | 2.96 | 3.92 |
| 4 | 6.64 | 66.51 | 20.25 | 3.41 | 3.19 |
| 6 | 9.43 | 59.43 | 25.12 | 3.78 | 2.23 |
| 8 | 11.59 | 53.32 | 28.12 | 5.80 | 1.17 |
| mean | | | | | |
| dpi | apoptotic | G0/1 | early S | S | G2/M |
| 0 | 0.45 | 90.81 | 2.09 | 3.73 | 2.92 |
| 2 | 2.27 | 73.25 | 18.86 | 3.30 | 2.32 |
| 4 | 5.88 | 68.64 | 17.62 | 4.23 | 3.63 |
| 6 | 8.67 | 64.07 | 18.95 | 4.44 | 3.87 |
| 8 | 13.20 | 60.81 | 19.26 | 4.26 | 2.48 |

Table S 3 Corresponding to figure 3: Cell cycle distribution of EBVwt infected B cells analyzed by PI staining. The mean was used to generate the plot in figure 3A. dpi – days post-infection

| **- CD40L feeder cells** | | | | |
|---|---|---|---|---|
| Donor 8 | | | | |
| dpi | apoptotic | G0/1 | S | G2/M |
| 0 | 0.14 | 97.93 | 1.44 | 0.49 |
| 2 | 5.85 | 85.99 | 5.34 | 2.82 |
| 4 | 9.66 | 70.71 | 10.82 | 8.80 |
| 6 | 12.57 | 68.51 | 10.64 | 8.28 |
| 8 | 17.71 | 66.19 | 9.05 | 7.05 |
| Donor 9 | | | | |
| dpi | apoptotic | G0/1 | S | G2/M |
| 0 | 0.18 | 97.90 | 1.43 | 0.49 |
| 2 | 9.49 | 82.18 | 5.55 | 2.78 |
| 4 | 17.93 | 70.52 | 6.89 | 4.65 |
| 6 | 12.65 | 68.01 | 10.53 | 8.81 |
| 8 | 19.34 | 64.30 | 8.59 | 7.78 |
| Donor 10 | | | | |
| dpi | apoptotic | G0/1 | S | G2/M |
| 0 | 0.21 | 97.69 | 1.63 | 0.47 |
| 2 | 5.65 | 86.13 | 5.39 | 2.83 |
| 4 | 9.68 | 70.95 | 10.93 | 8.44 |
| 6 | 12.84 | 68.23 | 10.92 | 8.02 |
| 8 | 17.61 | 65.99 | 9.08 | 7.32 |
| mean | | | | |
| dpi | apoptotic | G0/1 | S | G2/M |
| 0 | 0.18 | 97.84 | 1.50 | 0.48 |
| 2 | 7.00 | 84.77 | 5.43 | 2.81 |
| 4 | 12.43 | 70.73 | 9.54 | 7.30 |
| 6 | 12.69 | 68.25 | 10.69 | 8.37 |
| 8 | 18.22 | 65.49 | 8.90 | 7.38 |

| **+ CD40L feeder cells** | | | | |
|---|---|---|---|---|
| Donor 8 | | | | |
| dpi | apoptotic | G0/1 | S | G2/M |
| 0 | 0.14 | 97.93 | 1.44 | 0.49 |
| 2 | 7.55 | 73.50 | 12.03 | 6.91 |
| 4 | 6.91 | 67.54 | 16.81 | 8.74 |
| 6 | 18.03 | 63.22 | 11.55 | 7.20 |
| 8 | 29.79 | 55.41 | 7.67 | 7.13 |
| Donor 9 | | | | |
| dpi | apoptotic | G0/1 | S | G2/M |
| 0 | 0.18 | 97.90 | 1.43 | 0.49 |
| 2 | 6.91 | 72.73 | 13.06 | 7.30 |
| 4 | 11.40 | 60.67 | 15.59 | 12.34 |
| 6 | 15.55 | 64.93 | 11.81 | 7.70 |
| 8 | 30.47 | 55.65 | 7.11 | 6.77 |
| Donor 10 | | | | |
| dpi | apoptotic | G0/1 | S | G2/M |
| 0 | 0.21 | 97.69 | 1.63 | 0.47 |
| 2 | 11.80 | 76.43 | 8.47 | 3.30 |
| 4 | 7.05 | 67.22 | 18.02 | 7.71 |
| 6 | 17.73 | 63.01 | 11.14 | 8.12 |
| 8 | 30.24 | 55.80 | 7.66 | 6.30 |
| mean | | | | |
| dpi | apoptotic | G0/1 | S | G2/M |
| 0 | 0.18 | 97.84 | 1.50 | 0.48 |
| 2 | 8.76 | 74.23 | 11.18 | 5.83 |
| 4 | 8.42 | 65.19 | 16.82 | 9.57 |
| 6 | 17.10 | 63.72 | 11.50 | 7.68 |
| 8 | 30.17 | 55.62 | 7.48 | 6.73 |

Table S 4 Corresponding to figure 3 Cell cycle distribution EBVΔα1 infected B cells analyzed by PI staining. The mean was used to generate the plot in Figure 3A. dpi – days post-infection

| **- CD40L feeder cells** | | | | | | **+ CD40L feeder cells** | | | | |
|---|---|---|---|---|---|---|---|---|---|---|
| Donor 8 | | | | | | Donor 8 | | | | |
| dpi | apoptotic | G0/1 | S | G2/M | | dpi | apoptotic | G0/1 | S | G2/M |
| 0 | 0.14 | 97.93 | 1.44 | 0.49 | | 0 | 0.14 | 97.93 | 1.44 | 0.49 |
| 2 | 9.52 | 83.53 | 4.63 | 2.32 | | 2 | 10.44 | 58.55 | 19.96 | 11.05 |
| 4 | 14.69 | 75.36 | 5.12 | 4.83 | | 4 | 6.34 | 71.03 | 16.12 | 6.52 |
| 6 | 18.99 | 71.12 | 5.21 | 4.68 | | 6 | 13.78 | 73.30 | 10.06 | 2.87 |
| 8 | 20.44 | 71.90 | 4.08 | 3.58 | | 8 | 16.80 | 70.51 | 8.69 | 4.00 |
| Donor 9 | | | | | | Donor 9 | | | | |
| dpi | apoptotic | G0/1 | S | G2/M | | dpi | apoptotic | G0/1 | S | G2/M |
| 0 | 0.18 | 97.90 | 1.43 | 0.49 | | 0 | 0.18 | 97.90 | 1.43 | 0.49 |
| 2 | 10.45 | 82.20 | 4.80 | 2.54 | | 2 | 18.47 | 53.08 | 17.09 | 11.36 |
| 4 | 14.49 | 74.06 | 6.38 | 5.08 | | 4 | 7.52 | 68.80 | 14.21 | 9.47 |
| 6 | 15.66 | 73.16 | 5.58 | 5.60 | | 6 | 11.96 | 75.11 | 10.04 | 2.89 |
| 8 | 18.82 | 72.97 | 4.19 | 4.02 | | 8 | 17.88 | 69.79 | 8.66 | 3.68 |
| Donor 10 | | | | | | Donor 10 | | | | |
| dpi | apoptotic | G0/1 | S | G2/M | | dpi | apoptotic | G0/1 | S | G2/M |
| 0 | 0.21 | 97.69 | 1.63 | 0.47 | | 0 | 0.21 | 97.69 | 1.63 | 0.47 |
| 2 | 9.24 | 83.75 | 4.60 | 2.40 | | 2 | 9.57 | 59.33 | 19.54 | 11.56 |
| 4 | 15.14 | 74.50 | 5.73 | 4.62 | | 4 | 6.24 | 71.14 | 16.45 | 6.18 |
| 6 | 20.62 | 70.11 | 5.31 | 3.96 | | 6 | 13.78 | 73.41 | 10.04 | 2.78 |
| 8 | 21.32 | 71.00 | 3.97 | 3.71 | | 8 | 16.92 | 70.98 | 8.72 | 3.38 |
| mean | | | | | | mean | | | | |
| dpi | apoptotic | G0/1 | S | G2/M | | dpi | apoptotic | G0/1 | S | G2/M |
| 0 | 0.18 | 97.84 | 1.50 | 0.48 | | 0 | 0.18 | 97.84 | 1.50 | 0.48 |
| 2 | 9.74 | 83.16 | 4.68 | 2.42 | | 2 | 12.76 | 57.03 | 18.88 | 11.32 |
| 4 | 14.77 | 74.64 | 5.74 | 4.84 | | 4 | 6.70 | 70.33 | 15.59 | 7.38 |
| 6 | 18.43 | 71.46 | 5.37 | 4.74 | | 6 | 13.17 | 73.94 | 10.05 | 2.84 |
| 8 | 20.19 | 71.96 | 4.08 | 3.77 | | 8 | 17.20 | 70.43 | 8.69 | 3.69 |

Table S 5 Corresponding to figure 3 Cell cycle distribution of non-infected B cells analyzed by PI staining. The mean was used to generate the plot in figure 3A. dpi – days post-infection

| **- CD40L feeder cells** | | | | |
|---|---|---|---|---|
| Donor 8 | | | | |
| dpi | apoptotic | G0/1 | S | G2/M |
| 0 | 0.14 | 97.93 | 1.44 | 0.49 |
| 2 | 10.33 | 85.16 | 3.63 | 0.87 |
| 4 | 16.33 | 79.14 | 3.59 | 0.94 |
| 6 | 22.70 | 67.70 | 6.81 | 2.79 |
| 8 | 26.69 | 67.23 | 4.30 | 1.78 |
| Donor 9 | | | | |
| dpi | apoptotic | G0/1 | S | G2/M |
| 0 | 0.18 | 97.90 | 1.43 | 0.49 |
| 2 | 14.44 | 80.53 | 3.67 | 1.35 |
| 4 | 19.33 | 76.01 | 3.54 | 1.12 |
| 6 | 24.07 | 67.47 | 5.52 | 2.93 |
| 8 | 26.65 | 66.67 | 4.02 | 2.65 |
| Donor 10 | | | | |
| dpi | apoptotic | G0/1 | S | G2/M |
| 0 | 0.21 | 97.69 | 1.63 | 0.47 |
| 2 | 9.16 | 86.24 | 3.89 | 0.71 |
| 4 | 15.36 | 79.83 | 3.62 | 1.18 |
| 6 | 24.50 | 66.33 | 6.28 | 2.89 |
| 8 | 28.07 | 66.17 | 4.08 | 1.67 |
| mean | | | | |
| dpi | apoptotic | G0/1 | S | G2/M |
| 0 | 0.18 | 97.84 | 1.50 | 0.48 |
| 2 | 12.40 | 79.70 | 5.67 | 2.23 |
| 4 | 17.01 | 78.32 | 3.59 | 1.08 |
| 6 | 23.76 | 67.16 | 6.20 | 2.87 |
| 8 | 27.14 | 66.69 | 4.14 | 2.03 |

| **+ CD40L feeder cells** | | | | |
|---|---|---|---|---|
| Donor 8 | | | | |
| dpi | apoptotic | G0/1 | S | G2/M |
| 0 | 0.14 | 97.93 | 1.44 | 0.49 |
| 2 | 13.60 | 72.25 | 9.51 | 4.64 |
| 4 | 9.05 | 64.10 | 14.80 | 12.04 |
| 6 | 11.06 | 78.92 | 6.49 | 3.53 |
| 8 | 19.50 | 70.78 | 4.91 | 4.81 |
| Donor 9 | | | | |
| dpi | apoptotic | G0/1 | S | G2/M |
| 0 | 0.18 | 97.90 | 1.43 | 0.49 |
| 2 | 12.66 | 73.05 | 9.22 | 5.07 |
| 4 | 6.98 | 65.16 | 15.78 | 12.09 |
| 6 | 12.30 | 77.30 | 6.04 | 4.36 |
| 8 | 19.94 | 69.52 | 4.98 | 5.57 |
| Donor 10 | | | | |
| dpi | apoptotic | G0/1 | S | G2/M |
| 0 | 0.21 | 97.69 | 1.63 | 0.47 |
| 2 | 12.99 | 73.38 | 8.84 | 4.79 |
| 4 | 9.26 | 65.20 | 14.51 | 11.04 |
| 6 | 10.70 | 78.50 | 6.60 | 4.21 |
| 8 | 18.67 | 71.41 | 5.33 | 4.59 |
| mean | | | | |
| dpi | apoptotic | G0/1 | S | G2/M |
| 0 | 0.18 | 97.84 | 1.50 | 0.48 |
| 2 | 13.08 | 72.89 | 9.19 | 4.83 |
| 4 | 8.43 | 64.82 | 15.03 | 11.72 |
| 6 | 11.35 | 78.24 | 6.38 | 4.03 |
| 8 | 19.37 | 70.57 | 5.07 | 4.99 |
